# Red-light-only control of protein-protein interactions using a cyanobacteriochrome (UNICYCL)

**DOI:** 10.1101/2025.10.09.681461

**Authors:** Giang N.T Le, P. Maximilian M. Reed, Jaewan Jang, Kun Tang, Matias Zurbriggen, Maruti Uppalapati, Andrew Woolley

**Affiliations:** Department of Chemistry, University of Toronto, 80 St. George St., Toronto, ON, M5S 3H6, Canada; Inst. Of Synthetic Biology, Heinrich-Heine-Universität, Universitätsstr. 1, 40225 Düsseldorf, Germany; Department of Pathology and Laboratory Medicine, University of Saskatchewan, Saskatoon, Saskatchewan S7N 5E5, Canada

## Abstract

Most optogenetic tools are controlled by blue light. Red-light responsive tools enable multiwavelength applications and allow greater biological tissue penetration with reduced toxicity. Current red-light tools are primarily based on phytochromes, large dimeric proteins with a structurally complex mode of interaction with their binding partners. Here we introduce a small red-light-only responsive system composed of a GA domain (6 kDa) that binds to a CBCR GAF domain (17 kDa) with a K_d_ ∼1-5 µM to form a 1:1 complex in the dark. Red light causes dissociation of the complex by causing a >25-fold decrease in binding affinity. The CBCR GAF domain reverts to the dark state with a half-life of ∼1 min and the complex reforms. Structural analysis using NMR measurements combined with molecular docking and dynamics simulations shows that the binder interacts with the GAF domain and senses isomerization of the bilin chromophore at a site that overlaps the critical tongue domain of phytochromes. This system provides a small, simple red-light-only optogenetic tool that can operate to control protein-protein interactions *in vitro* and in living cells.

## Introduction

The use of light to cause protein-protein association or dissociation is the basis of many optogenetic tools that can be used to regulate biological processes with high spatiotemporal resolution.^1^ Most current optogenetic tools respond to blue light,^2^ however, red-light has lower levels of toxicity and better tissue penetration^3^ and enables orthogonal two-color control. The development of red-light responsive tools is therefore of wide interest.^4^

To date, red-light-responsive tools have been developed based on bacterial and plant phytochromes and their natural binding partners, PpsR2/Q-PAS1,^5^ phytochrome interacting factors (PIFs),^6^ or the *de novo* binders nanobody LDB3^7^ or affibody Aff6.^8^ However, phytochromes are large, dimeric proteins, features that complicate biophysical analysis as well as viral packaging and *in vivo* applications. There have been several attempts to reduce the size of phytochrome-based tools. For example, truncated PhyA, a red/far-red photoreceptor from *A. thaliana* and its binding partner FHY1 are small enough to be compatible with AAV vectors.^9^ Qiao et al improved the PhyA system for transcriptional control in higher organisms by fusing the N-terminal extension of PhyA to the photosensory module of the bacteriophytochrome *Dr*BphP.^10^ Red light dependent binding to the nanobody LDB3 allowed for the precise control of gene expression. Nevertheless, the smallest of these photosensory components still form dimers of mass >100 kDa. This is much larger than corresponding blue light tools like the *As*LOV domain (17 kDa).

The structural characterization of optogenetic systems can facilitate their effective application.^11^ For example, the structural characterization of LOV domains^12^ has enabled the design of blue light responsive transcription factors,^13^ Cas9,^14^ enzymes,^15^ and nuclear export/import sequences.^16^ In addition, structural characterization has enabled rational tuning of photocycle kinetics and the relative stabilities of on and off states.^17^ While significant progress has been made on the structural characterization of red-light absorbing bacteriophytochromes and plant phytochromes, how these interact with their binding partners to function as optogenetic tools has relied on model building^18^ until recently, when the structure of the Pfr state of PhyB bound to PIF6 was reported.^19, 20^ The PhyB/PIF6 structure reveals a large-scale reorganization of the PhyB dimer coupled to folding of the disordered PhyB N-terminal domain which then interacts with PIF6. One PIF6 molecule binds in an asymmetrical fashion to the dimer resulting in 1:2 PIF6:PhyB stoichiometry. The complexity of this interaction is reflected in the complex dynamics of photoswitching and thermal reversion.^20, 21^

Recently, we introduced cyanobacteriochrome (CBCR) cGMP-specific phosphodiesterases, adenylyl cyclases and FhlA (GAF) domains as components of red-light responsive tools, termed bidirectional, cyanobacteriochrome-based light-inducible dimers (BICYCLs).^22^ CBCR GAF domains can act as monomers and are much smaller than phytochromes (17 kDa vs. >100 kDa for the DrBphP photosensory domain). Using phage display, small binding partners (6 kDa) based on the GA domain scaffold can be selected that form 1:1 complexes with different photostates of CBCR GAF domains.^23, 24^ CBCR-GAF domain/binder pairs are therefore much simpler optogenetic tools than those based on phytochromes.

Here we introduce a CBCR-based red-light-only system that we denote UNICYCL. BICYCLs are bistable systems; red light converts the red-absorbing Pr state to the green-absorbing Pg state, and green light converts the Pg state back to the Pr state. Dual-wavelength optogenetic control is desirable for certain use cases, but not others. For instance, a bistable system can enable spatial patterning^22^ whereas a system that is activated by red light only and automatically turns off in the dark avoids the use of (less penetrating) green light and reduces the number of wavelengths required to control the system. CBCR-based tools that control cAMP synthesis have been specifically engineered with the goal of reducing thermal half-life to allow single-wavelength control.^25^

The UNICYCL consists of the red-light responsive CBCR GAF domain NpF2164g6^26^ and binders selective for the dark state of NpF2164g6. Photoisomerization of NpF2164g6 causes binder dissociation and can be used to reversibly colocalize proteins to agarose beads *in vitro*. The UNICYCL system can also function in mammalian cells to control gene expression using red light only, indicating a portability comparable to the BICYCL designs. In addition to these functional tests, we report a complete nuclear magnetic resonance (NMR) spectroscopy backbone triple-resonance assignment of NpF2164g6 in the dark state and an assignment of ∼80% of signals in the light state. We use these data to calculate a binding mode for the family of NpF2164g6 Pr-state binders we have developed. This analysis is expected to enable rational modification, tuning, and application of the UNICYCL system.

## Experimental Methods

### NpF2164g6 Expression and Preparation for Phage Display

The NpF2164g6 (1038-1206) gene was a gift from J. Clark Lagarias. It was cloned into a version of the pBAD His B plasmid that included an N-terminal Avi tag and a TEV cleavage site via Gibson assembly. Another version of NpF2164g6 was made in this plasmid by removal of the TEV site via Gibson assembly. *E. coli* strain BL21(AI) (Thermo-Fisher) was co-transformed with this plasmid and pPL-PCB (a gift from J. Clark Lagarias.^27^ Transformed cells in 100 mL of LB media (supplemented with 50 µg/mL kanamycin, 100 µg/mL ampicillin) were grown at 37 °C until an OD of 0.5 was reached. The cells were transferred into 1 L of LB (50 µg/mL kanamycin, 100 µg/mL ampicillin, 1 mM IPTG) and grown for 1 h, after which 0.02% arabinose was added for NpF2164g6 induction, and the culture was incubated further for 1.5 h at 37 °C. The temperature was lowered to 20 °C, and cells were grown for 12-18 hours. The cells were centrifuged and resuspended in 50 mM phosphate buffer with 300 mM NaCl at pH 7.5. Lysis was then conducted by sonication and cells were centrifuged at 10K RPM for 1 hour. The supernatant was filtered and then loaded onto Ni-NTA beads (Thermo Fisher) to purify holo-NpF2164g6 protein. Proteins were subsequently eluted with lysis buffer containing 250 mM imidazole. Eluted proteins were dialyzed into PBS (10 mM Na_2_HPO4, 1.8 mM KH_2_PO_4_, 137 mM NaCl, 2.7 mM KCl, pH 7.2) and further purified via size exclusion chromatography (using a Superdex 75 10/300 GL column (GE Healthcare) running at a flow rate of 0.4 mL/min).

Biotinylated-NpF2164g6 was prepared according to a published protocol.^28^ The reaction mixture of 100 µM Avi-tagged holo-protein in 5 mM MgOAc, 2 mM ATP, 1 µM BirA enzyme, and 150 µM D-biotin was incubated for 1 hour at room temperature followed by overnight incubation at 4 °C. Biotinylated holo-NpF2164g6 was further purified by size exclusion chromatography as described above and its mass was confirmed via electrospray ionization mass spectrometry (ESI-MS). For long-term storage, biotinylated protein in storage buffer (10 mM Na_2_HPO_4_, 1.8 mM KH_2_PO_4_, 137 mM NaCl, 2.7 mM KCl, 10% glycerol, pH 7.2), flash frozen using liquid nitrogen and stored at −80 ◦C.

### Phage Display Binder Expression

Hits obtained from phage display were cloned into the pET24 plasmid with C-terminal poly His (6x) tags via Gibson assembly. Proteins were expressed by growth in LB at 37 °C until OD600∼0.6, followed by addition of IPTG to 1 mM, lowering of temperature to 18 °C, and growth under those conditions for 12-18 hours. Cells were lysed by resuspension in denaturing buffer (6M guanidinium HCl, 100 mM phosphate, 10 mM imidazole, pH 8.0) followed by sonication. Cells were centrifuged and the supernatant was filtered and then applied to a Ni-NTA column as described above. Protein was eluted from the column with denaturing elution buffer (6M guanidinium HCl, 100 mM acetate, 10 mM imidazole, pH 4.5) then dialyzed into PBS (pH 7.2). Proteins were then further purified by size exclusion chromatography as described above.

### NpF2164g6 Expression for NMR Spectroscopy

Standard protein NMR relies on labeling of proteins with ^15^N and ^13^C from labeled ammonium chloride and glucose respectively. However, since glucose interferes with the activity of the pBAD promoter, we used a different plasmid for expression of isotopically labeled NpF2164g6. NpF2164g6 (1038-1206) was cloned via Gibson assembly into the pMH1105 plasmid,^29^ a kind gift from Wilfried Weber. This plasmid enables the use of IPTG as the inducer for expression of a protein-of-interest alongside the HO1 and PcyA genes necessary for PCB synthesis in *E. coli*. A version of NpF2164g6 containing an N-terminal truncation, NpF2164g6 (1052-1206), was also cloned into pMH1105, and was the version used for triple resonance assignment. For labeled protein expression, cells were grown in M9 media supplemented with: 1 g/L NH_4_Cl, 2 g/L glucose, 0.1 mM CaCl_2_, 1 mM MgSO_4_, 25 µM ZnSO_4_, 50 µM FeCl_3_ (made day-of), 100 mg/L streptomycin, 5 mg/L biotin, 5 mg/L thiamine, 1 mg/L of each of choline chloride, folic acid, niacinamide, D-pantothenate, and pyroxidol, and 0.1 mg/L of riboflavin. pMH1105 with NpF2164g6 was transformed into BL21 (DE3) *E. coli*, inoculated in LB, and grown overnight. The overnight culture was centrifuged and the cells transferred to a 50 mL unlabeled M9 starter culture (initial OD600 ∼0.3). The starter culture was grown at 37 °C and 180 RPM to an OD600 of ∼0.6, then the cells were centrifuged at 4K RPM for 10 minutes. The pellet was resuspended and then transferred to 1 L of labeled M9 media. The cells were grown to an OD600 of 0.8, then IPTG was added to 1 mM and the temperature was changed to 18 °C. The cells were then grown overnight (12-18 hours). This procedure was also followed to produce proteins for thermal reversion and fluorescence titrations. Proteins were purified as described above, with the exception that cell lysis was carried out for labeled protein samples by first flash-freezing and thawing cells 5 times, then passing cells through a homogenizer EmulsiFlex-C3 high pressure homogenizer to assure complete lysis.

### Phage Display

We used a previously established phage display protocol^22-24^ to generate state-specific binders for NpF2164g6. Biotinylated NpF2164g6 was immobilized on streptavidin-coated 96-well plates and either irradiated with 660 nm light or left in the dark during selection. The naïve GA domain library was displayed on the pVIII coat protein of M13 phage. The library was subjected to three rounds of selections to identify the initial hits. The most promising clone, BNp-Red-1.0, showed ∼40-fold selection to the Pr state versus the Pg state. To increase the affinity of BNp-Red-1.0 toward the Pr-state, we conducted a second selection with a new library based on BNp-Red-1.0 (soft-randomization approach) displayed on the pIII coat protein. The following oligonucleotide was used for mutagenesis:

5’-<u>AAGGCTGGTATCACC</u>(N4)(N2)(N4)GAC(N3)(N2)(N3)(N4)(N1)(N4)TTCAAC(N4)(N4)(N1)ATC AAT(N3)(N1)(N4)GCG(N4)(N4)(N4)(N3)(N1)(N4)GTG(N3)(N1)(N4)(N4)(N1)(N4)GTTAAC(N4)(N1) (N4)(N2)(N3)(N3)AAGAAC(N1)(N1)(N3)<u>ATCCTGAAAGCTCAC</u>-3’

Where N1 is a mix of 70% A, 10% C,10% G, 10% T

N2 is a mix of 10% A, 70% C, 10% G, 10% T

N3 is a mix of 10% A, 10% C, 70% G, 10% T

N4 is a mix of 10% A, 10% C, 10% G, 70% T

This library underwent three rounds of selection in which a final light-elution step was introduced^23^ to obtain variants with improved affinity and dynamic range.

### Size Exclusion Chromatography

Purified protein samples in PBS (pH 7.2) were injected onto a Superdex 75 10/300 GL column (GE Healthcare) equilibrated with PBS (pH 7.2). The column flow rate was set to 0.4 mL/min. Samples were either kept in darkness or irradiated with a custom LED array of Endor Star 7040 LUXDRIVE Red LEDs. All proteins were injected at a concentration of 30 µM and a volume of 0.5 mL. The column was calibrated with a standard composed of: aprotinin (6.5 kDa), ribonuclease A (13.7 kDa), carbonic anhydrase (29 kDa), and ovalbumin (43 kDa).

### UV-Vis and Fluorescence Characterization

The concentration of NpF2164g6 was quantified by dilution of the protein into 6M guanidinium HCl at pH 1 under dark conditions. The concentration was determined from absorbance using ϵ_662nm_ = 35500 M^−1^ cm^−1^.^30^ Binder concentrations were determined based on A_280nm_ readings on an Implen NanoPhotometer N60. UV-Vis spectra were taken on a BMG Labtech SPECTROstar Nano. Fluorescence readings and UV-Vis kinetic traces were taken on a BioTek Synergy H1 microplate reader. All readings were taken at 25 °C in pH 7.2 PBS (recipe above).

### NMR Spectroscopy

All spectra were acquired at 20 °C in pH 7.2 PBS. Spectra were acquired either at the University of Toronto Department of Chemistry CSICOMP NMR Facility on a 700 MHz Agilent DD2 NMR spectrometer with a cryogenically cooled probe (dark and light ^1^H-^15^N HSQCs, dark and light HNCO, dark and light HNCA, dark HNCACB, and dark HN(CO)CACB) or the University of Toronto Biochemistry Nuclear Magnetic Resonance Centre on a Varian 600 MHz spectrometer with a triple resonance cryo-probe (HN(CA)CO (dark) as well as a dark ^1^H-^15^N HSQC to assure the result was the same as on the 700 MHz spectrometer). All chemical shifts reported are those obtained with the 700 MHz spectrometer. Pg state spectra were acquired by inserting a fiber optic cable into the sample tube and irradiating with a Thorlabs M660L3 LED while spectrum acquisition proceeded. The BioPack watergate 15N-HSQC pulse sequence was used to obtain all ^1^H-^15^N HSQC spectra. All other spectra were also obtained using the BioPack versions of the pulse sequences, except for the HN(CA)CO which used the the sequence described by Clubb *et al*.^31^ NMR samples were made to a volume of ∼400 µL with ∼15% D_2_O. All 3D spectra were acquired on NpF2164g6 (1052-1206), while ^1^H-^15^N HSQC spectra were acquired for both NpF2164g6 (1038-1206) and NpF2164g6 (1052-1206).

To categorize what constituted an affected residue when analyzing HSQC spectra of NpF2164g6 with and without BNp-Red-1.0, we developed a more systematic version of our previous protocol. ^32^ The sample contained 120 µM ^15^N-labeled truncated NpF2164g6 and 240 µM BNp-Red-1.0. All peaks that were no longer visible at a low contour level after BNp-Red-1.0 addition were counted as affected. Following this, remaining assignments were transferred from the unbound to bound NpF2164g6 peaks. When assignment transfer was ambiguous, all residues involved were excluded from analysis. For each assigned residue, the ratio (intensity with BNp-Red-1.0) / (intensity alone) was calculated alongside a change in chemical shift metric equal to 5|δH| + |δN|. The average change in intensity was divided by 2, and any residue with a change in intensity below this was counted as affected. The average chemical shift metric was multiplied by 2, and any residue with a change in chemical shift above this was counted as affected.

### Molecular Dynamics and Docking

Molecular dynamics (MD) protocols were identical to those described previously^33^ employing the Amber GAFF2 and FF14SB force fields with TIP3P water^34, 35^ and a PCB parametrization provided by Igor Schapiro.^36^ The engine used to conduct simulations was the GPU implementation of AMBER18.^37^ The only significant deviation from the prior procedure was that in this study no unrestrained equilibration step was included and, following restrained equilibration, the simulation immediately proceeded to unrestrained production MD for 1 µs.

The structure for NpF2164g6 (1052-1206) without PCB was generated by the ColabFold implementation of AlphaFold2.^38^ The top-ranked structure was aligned against the 3W2Z structure of AnPixJg2^39^ and the PCB chromophore was transferred from this structure and combined with residue C90 to create a single unnatural amino acid. Analysis of molecular dynamics trajectories was conducted in CPPTRAJ.^40^

We used the HADDOCK2.4 web server^41^ to generate structures of BNp-Red-1.0 bound to truncated NpF2164g6. We used ColabFold to generate a structure of BNp-Red-1.0 for input to the program.^38^ The active residues of NpF2164g6 (1052-1206) were set to be all those affected by BNp-Red-1.0 addition, and the active residues of BNp-Red-1.0 were set to be those mutated during phage display. All settings were at default values. After the initial MD run of the top-ranked structures from each of the top two-ranked HADDOCK clusters, a representative frame from each run was selected for further simulation in replicate. To do this, all frames within a trajectory were aligned against all other frames of that trajectory using the non-terminal residues of NpF2164g6. This resulted in frames where NpF2164g6 is in an essentially fixed position while BNp-Red-1.0 moves freely. Within each trajectory, a relatively stable portion was determined by inspection of the Cα RMSD against the initial frame. A representative frame was then extracted from this segment, using the following metric: after alignment of the complexes by the NpF2164g6 non-terminal residues, we denote the displacement of the alpha carbon of an BNp-Red-1.0 residue n between frame i and frame j as 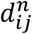. A metric of global similarity within the stable region was computed for each frame i equal to 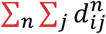.The representative frame was chosen to be the frame with the lowest value for this metric.

### SEAP Assay

The Sleeping Beauty (SB) SB100X transposase system was used for genome engineering^42, 43^ NpF2164g6 and the binder A12 were cloned into the plasmid pKT1308 (ITR-P_EF1a_-NpF2164g6-FUS-VP16-NLS-IRES-E-A12-NLS, P_RPBSA_-Puro^R^-pA-ITR) together with a SEAP reporter pDD123 (ITR-etr8-Pmin-SEAP-pA, PRPBSA-HygR-pA-ITR)^44^ for stable integration into the genome. A total of 200,000 CHO-K1 cells were seeded in a 6-well plate one day prior to transfection. The plasmid (1.8 µg) was co-transfected into CHO-K1 cells with 0.75 µg of the SB transposase expression vector (a gift from Zsuzsanna Izsvak (Addgene plasmid # 34879; http://n2t.net/addgene:34879; RRID:Addgene_34879))^42^ followed by selection with 10 µg mL^−1^ puromycin (Thermo-Fisher) and 400 *µ*g mL^*−*1^ hygromycin (InvivoGen). Afterwards, 96 cell clones were randomly selected and 10,000 cells were seeded into a 96 well plate. After 24 h, the cell culture media was changed with 40 µM PCB chromophore and illuminated for 24 h with 10 µmol m^−2^ s^−1^ light of 660 nm or kept in darkness. SEAP activity was analyzed as described previously.^22^

### Colocalization Assay

Both the mCherry-A12 and A12-mCherry fusion proteins were expressed and purified with a C-terminal poly His (6x) tag. Streptavidin-coated agarose beads (Sigma) were equilibrated by washing three times with HEPES buffer (10 mM HEPES, 150 mM NaCl, 1 mM EDTA, pH 7.4). To label the resin beads, either biotinylated NpF2164g6 or biotinylated IgG (Rockland) was incubated with the bead slurry for 1 hour in the dark. Excess unbound protein was removed by washing with HEPES buffer. One microliter of NpF2164g6-coated beads and 1 µL of IgG-coated beads were mixed with 50 µL of either 500 nM mCherry-A12 or A12-mCherry in HEPES buffer. The mixtures were incubated for 1 hour in the dark and then transferred to a 0.17-mm Greiner 96-well flat-bottom microplate for imaging. Bead imaging was performed using a Leica Mica microscope equipped with a built-in LED light source (555 nm, 170 mW; 625 nm, 170 mW) and a 20× air objective lens. mCherry fluorescence was captured using 555 nm excitation (9.6% intensity, 250 ms exposure). Photoconversion of NpF2164g6 was induced using 625 nm illumination (3.8% intensity). Image analysis was conducted in ImageJ using the Time Series Analyzer V3 plugin. Background subtraction was applied using the sliding paraboloid method to generate the final movie.

## Results & Discussion

### Development of a red-light-only optogenetic system based on NpF2164g6

We chose the red/green CBCR GAF domain NpF2164g6^26^ as the photoswitch for the UNICYCL system (Fig. 1). NpF2164g6 binds to the phycocyanobilin (PCB) chromophore. The thermally stable Pr state displays an absorbance peak at 647 nm (Fig. 1c); irradiation with red light produces the Pg state with an absorbance peak at 550 nm.^33^ The Pg state thermally reverts to the Pr state with a half-life of ∼1 minute (Fig. S1). In addition to its short thermal half-life, the NpF2164g6 protein has a high quantum yield (32% ^45^) compared to phytochromes (2-18%^46^) a useful feature for deep tissue applications where attenuation of light intensity is a significant problem.

**Figure 1:**
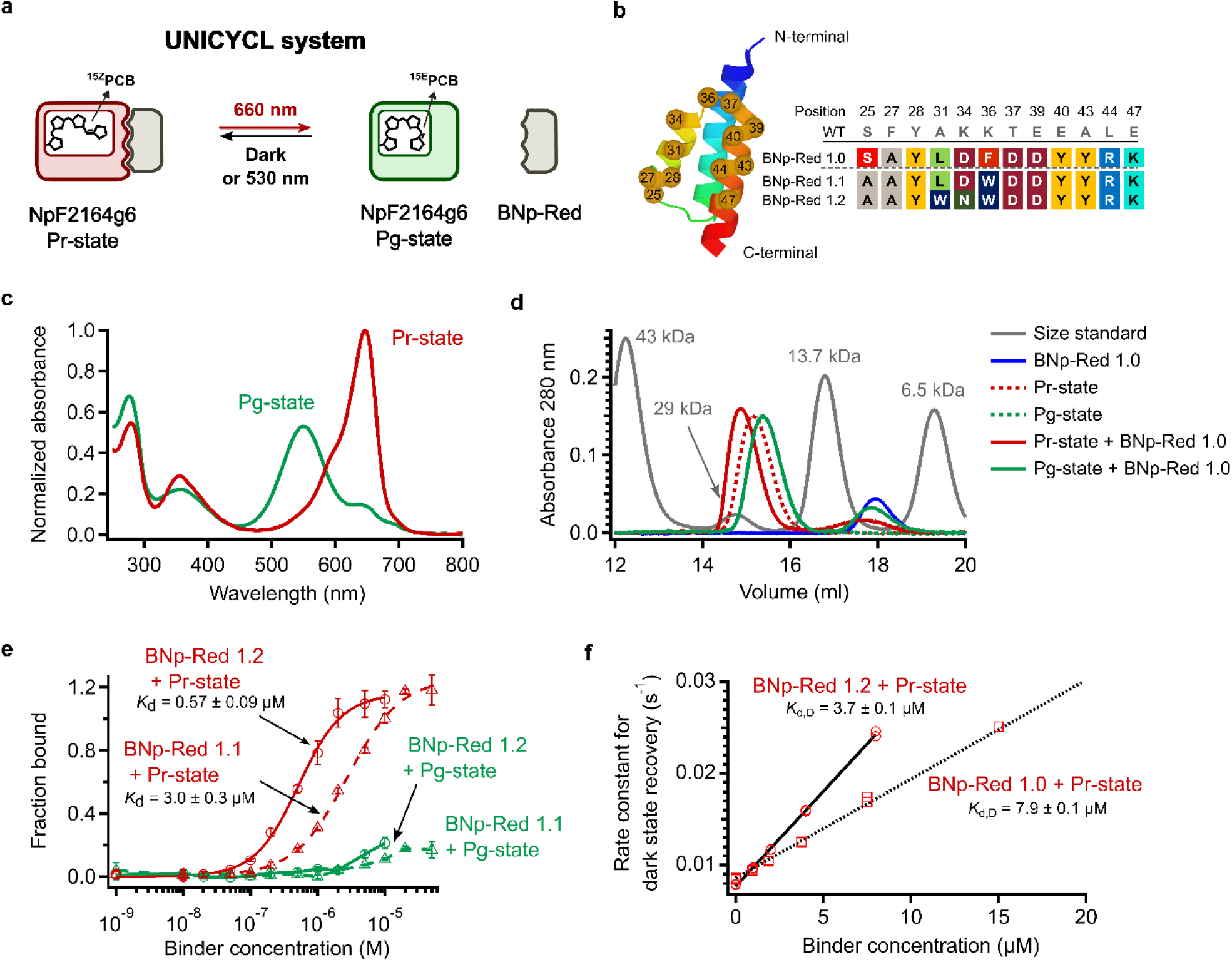
Development and characterization of NpF2164g6 binders. (a) Schematic illustration of the UNICYCL system. (b) 12 amino acids in the second and third helices of the GA domain (PDB: 1TF0) were randomized to generate a variable surface (left). Sequences of the randomized residues of BNp-Red-1.1-BNp-Red-1.2 binders (right). (c) UV-Vis spectra of the NpF2164g6 protein in its Pr (dark) state and Pg (light) state. The latter was obtained by irradiation with 660 nm light. (d) Size exclusion chromatography. Proteins were injected at 30 µM in all cases. NpF2164g6 and BNp-Red-1.0 show stronger association in the dark. The dash green trace is obscured because it is overlapped with the bold green trace. (e) ELISAs using purified biotinylated NpF2164g6 with purified BNp-Red-1.2 and BNp-Red-1.1. n = 4 technical replicates, mean ± sd. (f) Rate constant for thermal reversion of NpF2164g6 versus the concentration of BNp-Red-1.0 and BNp-Red-1.2. These lines can be fit to yield the dissociation constant for the dark state interaction, K_d,D_, on the assumption that the interactions have ϕ_switching_ values^32^ of 1 and the concentration of binder is much smaller than K_d,L_. n = 2 technical replicates.

To create a uni-wavelength, cyanobacteriochrome-based red-light responsive optogenetic tool we performed phage display against immobilized NpF2164g6 using the protocol we previously developed (see Methods). Naïve selection using a GA domain library produced two clones that displayed selective binding to the Pr state of NpF2164g6. These two clones were sequenced and found to be identical (Fig. 1b). The sequence, designated BNp-Red-1.0 (for ‘binder of NpF2164g6-Red state’), was cloned into a pET24 vector, then expressed and purified (see Methods). Figure 1d shows the size exclusion chromatography elution profile for purified NpF2164g6 in the dark-adapted Pr state and the Pg state (under constant red-light illumination). These species elute at slightly different volumes, both consistent with a monomeric protein of mass ∼17 kDa. The addition of an equimolar concentration of purified BNp-Red-1.0 to dark-adapted NPF2164G6 produced a species that eluted earlier, consistent with 1:1 NpF2164g6:BNp-Red-1.0 complex with a molecular weight of ∼25 kDa. Under red light, no complex could be detected (Fig. 1d).

We then performed affinity maturation using the BNp-Red-1.0 sequence as a template (see Methods) to find clones with greater Pr state selectivity and greater binding affinity to NpF2164g6. The phage-display-based selection procedure was modified by introducing a light-elution step to increase the stringency of the selection. Screening produced several hits, with clones BNp-Red-1.1 and BNp-Red-1.2 exhibiting the largest Pr to Pg signal ratio in phage ELISA assays, 26-fold and 30-fold selectivity, respectively (Fig. S2). These sequences (Fig. 1b) were found to be slight variations of BNp-Red-1.0. Clone BNp-Red-1.1 had the following mutations, Phe36Trp and Ser25Ala. Clone BNp-Red-1.2 had Leu31Phe and Asp34Asn when compared to BNp-Red-1.0. The BNp-Red-1.1 and BNp-Red-1.2 sequences were expressed with C-terminal FLAG tags to permit detection of binding to immobilized NpF2164g6 (see Methods). Figure 1e shows ELISAs using purified binders under dark-adapted and red-light conditions. Binding affinities for the dark state (Pr) were estimated to be 0.6 ± 0.1 µM for BNp-Red-1.2 and 3 ± 0.3 µM for BNp-Red-1.1. Binding to the red light irradiated state (Pg) was too weak to be reliably determined (>100 µM). These data indicate that binding was highly selective, *i*.*e*. > 25-fold tighter for the Pr state. (Fig. 1e).

### Thermal reversion and fluorescence

To determine binder affinities in solution we carried out measurements of NpF2164g6 thermal reversion rates as a function of binder concentration. Previously, we reported that binding proteins selective for the light or dark state often alter the thermal reversion rate of a photoswitchable protein.^32^ For CBCR GAF domains specifically, it was found that the change in the thermal reversion rate upon adding excess binder is equal to the ratio of the K_d_ values for the binder and the dark-state protein (NpF2164g6 Pr) and the binder and the light-state protein (NpF2164g6 Pg). That is, k_b_/k_u_ = K_d,L_/K_d,D_, where k_u_ is the rate constant for thermal reversion for the photoswitchable protein alone, k_b_ is the rate constant for thermal reversion of the bound protein, K_d,L_ is the dissociation constant of the binder from the light state, and K_d,D_ is the dissociation constant of the binder in the dark. Measuring the observed thermal reversion rate versus binder concentration allows determination of K_d,L_. Figure S3 shows these data; for the NpF2164g6 Pg state/BNp-Red-1.0 interaction a K_d,L_ greater than 200 µM was determined. For the NpF2164g6 Pg state/BNp-Red-1.2 interaction a K_d,L_ greater than 30 µM was determined.

As with other CBCR GAF domains,^22, 32^ fluorescence of the dark state of NpF2164g6 was found to be affected by binders. Figure S4 shows measured Pr state fluorescence as a function of BNp-Red-1.0 or BNp-Red-1.2 concentration. Fitting these data to a 1:1 binding model gives K_d,D_ values of 7 ± 3 and 3 ± 3 µM respectively. Additionally, as shown in Reed et al.^32^ when k_b_/k_u_ = K_d,L_/K_d,D_, then K_d,D_ can be determined using the equation k_obs_ = k_u_(1+B_tot_/ K_d,D_), where B_tot_ is total binder concentration, provided that B_tot_ << K_d,L_. The K_d,D_ values derived in this way agree well with the fluorescence titration values and have a much lower error (Fig. 1f, S4). These data confirm that the binders are highly selective, with a > 25-fold change in K_d_ upon irradiation for BNp-Red-1.0.

### The UNICYCL system functions in an in vitro colocalization assay

To determine whether the association of the UNICYCL system is reversible and controllable in a microscopy-based assay, we coated agarose bead surfaces with either NpF2164g6 or an unrelated protein (IgG) as a control and incubated the bead mixture in a solution containing 500 nM BNp-Red-1.2 fused to an N-terminal mCherry tag (Fig. 2a). In the dark state, mCherry-BNp-Red-1.2 localized specifically to NpF2164g6-coated beads, with no detectable binding to IgG-coated beads (Fig. 2b). Upon red light illumination, mCherry-BNp-Red-1.2 rapidly dissociated from the bead surface, indicating a light-dependent, reversible interaction. The association in the dark and dissociation under red light were observed over at least two cycles (Fig. 2b, Movie S1). Notably, when BNp-Red-1.2 was tagged with mCherry at the C-terminus, no bead localization was observed (Figure S5). This suggests that the C-terminal tag may sterically hinder the interaction interface of BNp-Red-1.2, thereby disrupting its association within the UNICYCL system.

**Figure 2:**
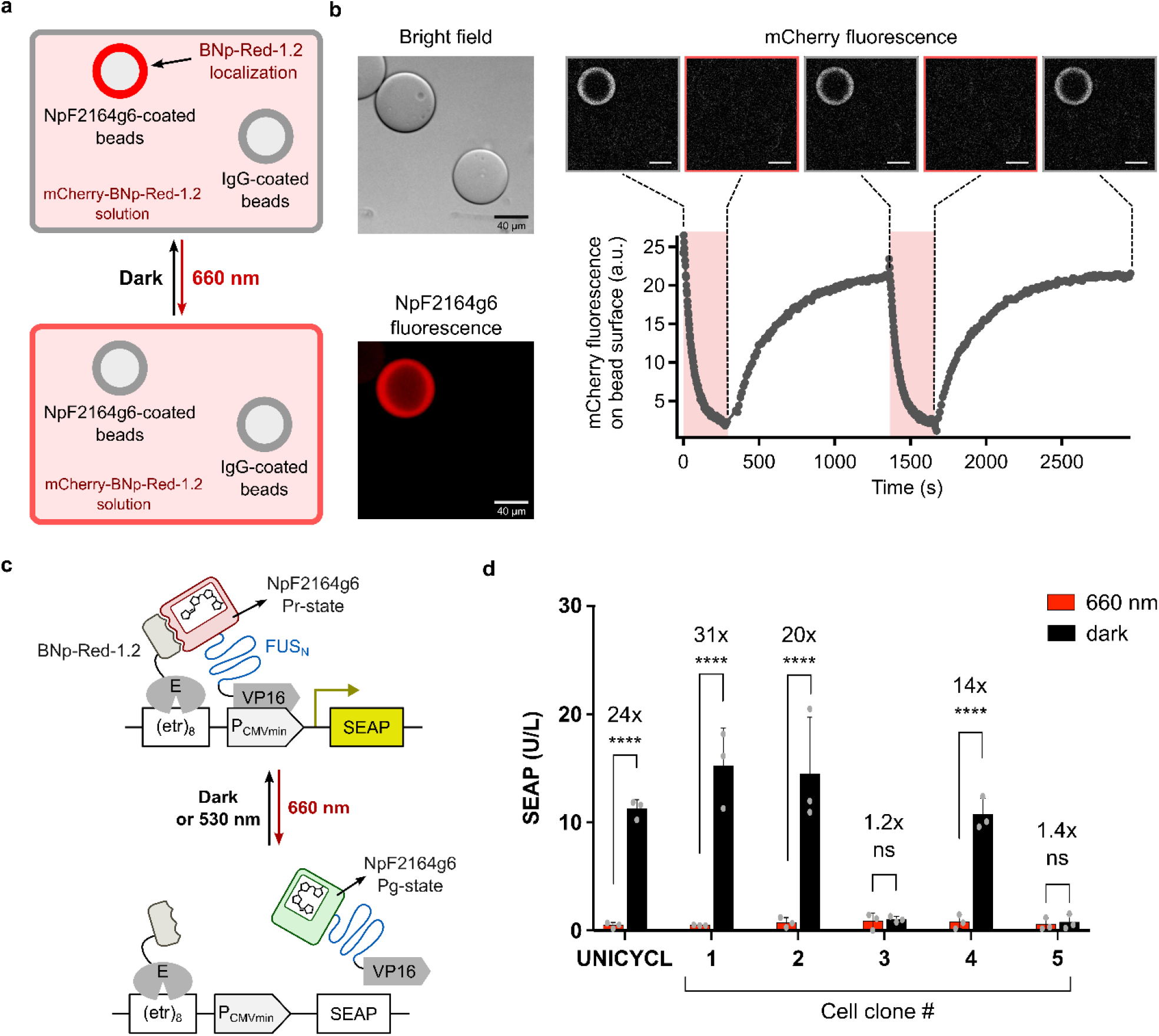
Application of the UNICYCL system *in vitro* and in mammalian cells. (a) Schematic diagram of light-induced protein localization on bead surfaces using the UNICYCL system. A mixture of NpF2164g6-coated and IgG-coated (negative control) agarose beads were resuspended in a solution containing 500 nM BNp-Red-1.2-mCherry. NpF2164g6-coated beads were identified using NpF2164g6 fluorescence. BNp-Red-1.2-mCherry localized on the surface of NpF2164g6-coated beads in the dark and completely dissociated into solution under red light illumination. (b) Left panel: bright-field image (top) and NpF2164g6 fluorescence image (bottom) of NpF2164g6-coated and IgG-coated beads. Bottom right panel: quantification of mCherry fluorescence intensity on bead surfaces during cycles of red-light illumination and darkness. Top right panel: widefield mCherry fluorescent images of beads at each time point. Images are representative of two replicate experiments. (c) Scheme of the UNICYCL controlled SEAP transcriptional reporter assay (d) Profile of randomly selected clones derived from the UNICYCL bulk culture. Data points represent SEAP values determined from a single sample for each clone measured with three replicates (the bar indicates the average).

### The UNICYCL system functions to control gene expression in mammalian cells

To assess the transferability of the UNICYCL system, we conducted an assay to test the ability of the system to control expression of a gene of interest in CHO-K1 cells. The alkaline phosphatase enzyme was placed under the control of the minimal promoter, (etr8)-PCMV min. NpF2164g6 was expressed in cells as a fusion with the VP16 promoter transactivator, while BNp-Red-1.2 was fused to the E (erythromycin repressor) DNA-binding protein (see Methods). The construct pKT1308 (EF1a-g6-FUS-VP16-NLS-IRES-E-BNp-Red-1.2-NLS) was stably integrated together with a SEAP reporter in CHO-K1 cells. After supplementing the cellular media with PCB chromophore, SEAP activity was measured under dark conditions vs. under 660 nm illumination. The bulk cell culture show ∼24-fold induction of gene expression (Fig. 2d); individual clones were in some cases non-switchable, whereas other showed up to >30-fold photoswitchability (Fig. 2d) likely reflecting the effects of integration site on expression of the UNICYCL system.^22, 47^

### Structural analysis of the UNICYCL system

Structural characterization of optogenetic tools can enable rational engineering to optimize behaviour.^11, 48^ We used NMR spectroscopy to characterize both the structural changes of NpF2164g6 upon photoswitching as well as the mode of binding of BNp-Red-1.0 and its derivatives. Our analyses are facilitated by the work of Ames, Lagarias and colleagues, who used NMR to perform a detailed characterization of the CBCR GAF domain NpR6012g4, which is highly homologous to NpF2164g6 (Fig. S6).^49^ We previously transferred assignments from NpR6012g4 to NpR6012g4 T631G for both the light and dark states.^33^ This set of assignments was then compared to those of NpF2164g6, which also has Gly in the W(S/G)GE motif.^33^

The NpF2164g6 construct used to generate binders, NpF2164g6 (1038-1206), contains an N-terminal region (residues 1038-1051) that is predicted by AlphaFold3 (with low confidence) to form an extension of the N-terminal helix (Fig. S7). In contrast, the crystal structure of the homologous CBCR GAF domain AnPixJg2^39^ (PDB code: 3W2Z) shows the corresponding region does not extend the N-terminal helix but instead turns and packs against the N- and C-terminal helices (Fig. S7). NMR HSQC spectra of NpF2164g6 and a variant with the N-terminal region removed (NpF2164g6 (1052-1206)) showed the same number of peaks outside of the center of the proton dimension (Fig. S8). This result implies the truncated region is not stably folded in solution but is undergoing conformational transitions that result in loss of NMR signal. Thermal reversion and fluorescence titrations of truncated NpF2164g6 with BNp-Red-1.0 showed similar photoswitchable binding (Fig. S9) and the truncated variant had a slightly lower k_u_ (Fig. S9). Based on the small effect of truncation and the lack of NMR signals associated with the N-terminal extension, we selected the truncated NpF2164g6 variant for further biophysical study.

### Structures of NpF2164g6, BNp-Red-1.0 and the NpF2164g6:BNp-Red-1.0 complex

Using a standard suite of triple-resonance experiments, we obtained a full backbone assignment of NpF2164g6 (1052-1206) in the dark (Pr) state using CA, CB, and CO connectivity. We also transferred 80% of assignments to a light state spectrum, with transfer of assignments aided by reference to the HNCA spectrum of the light (Pg) state (Fig. S10). Comparison of the changes in H and N chemical shifts upon irradiation in NpR6012g4 T631G to those in NpF2164g6 indicates that these proteins undergo very similar structural changes upon irradiation even though the former has a light state half-life of ∼2 hours, and the latter of ∼1 minute (Fig. S11).^33^

We constructed a model of the dark state of NpF2164g6 using ColabFold with default parameters.^38^ Predicted local distance difference test (pLDDT) scores^50^ were above 90 for most of the sequence except for the extend loop (residues 50-62; pLDDT ∼80-90) (*vide infra*), the chromophore attachment site (Fig. S12), and the extreme N- and C-termini. TALOSn predictions of protein secondary structure from the measured NMR chemical shifts^51^ are consistent with this predicted structure (Fig. S13). The PCB chromophore in its C5-*Z,syn* C10-*Z,syn* C15-*Z,anti* configuration was then linked to Cys89 resulting in *S* chirality at the C3^1^ atom as in NpR6012g4 (see MD methods).^49^ To confirm the model represented a stable structure for dark-adapted NpF2164g6 in solution, a 1 µs molecular dynamics (MD) simulation was performed in triplicate using the FF14SB force field (see MD methods). The calculated α-carbon RMSD was stable at ∼1 Å over the course of these trajectories (Fig. S14) confirming the model represented a stable structure.^52^

We used ColabFold to generate a structure of the binder BNp-Red-1.0. This structure was essentially the same as the wild-type GA domain (PDB code:1TF0) as expected. The predicted local distance difference test (pLDDT) scores were above 90 for the entire sequence (Fig. S15).

We then acquired an ^1^H-^15^N HSQC spectrum of NpF2164g6 in the presence of the parental binder BNp-Red-1.0. Upon addition of BNp-Red-1.0, peaks for several NpF2164g6 residues shift, others decrease in intensity, and others vanish entirely (Fig. 3, S10). These observations are similar to those reported in our previous work characterizing a photostate-selective binder to a CBCR.^32^ Based on the spectral changes seen, a set of residues affected by BNp-Red-1.0 addition was identified (see NMR Methods). Mapping the residues affected by BNp-Red-1.0 onto the dark state NpF2164g6 structure reveals a patch near the chromophore (Fig. 3). The residues affected by BNp-Red-1.0 binding have strong overlap with the residues affected by photoswitching (Fig. 3a,b, S10).

**Figure 3:**
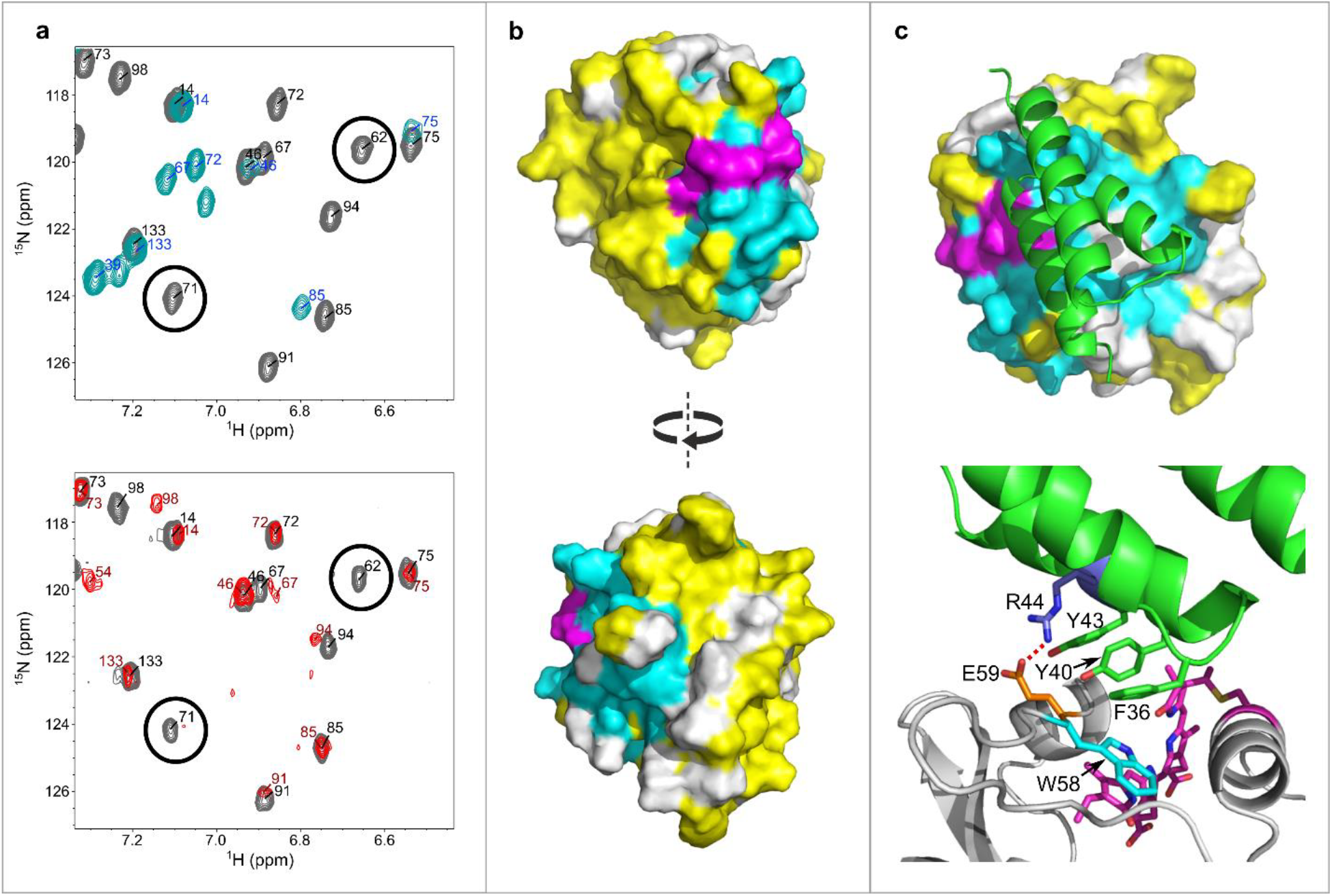
Structural characterization of NpF2164g6 (truncated variant). (a) (upper) ^1^H-^15^N TROSY HSQC spectra of NpF2164g6 in the dark (grey) or under constant irradiation with 660 nm light (cyan) with assignments marked in black and blue respectively. (lower) ^1^H-^15^N TROSY HSQC spectra of NpF2164g6 in the dark alone (grey) or with excess BNp-Red-1.0 (red) with assignments marked in black and red respectively. Two residues (62 and 71) affected similarly by light or by BNp-Red-1.0 addition are indicated by black circles. (b) NpF2164g6 shown as a surface. Cyan residues are affected by BNp-Red-1.0 addition, yellow residues are unaffected, and white residues have no data available. The chromophore is shown in magenta. (c) The modeled structure of BNp-Red-1.0 (green) bound to dark-state NpF2164g6. (lower)

We attempted to use ColabFold^38^ and AlphaFold3^53^ to generate models of BNp-Red-1.0-derived binders docked onto NpF2164g6, however, the models generated had no significant degree of agreement with the NMR data (Fig. S16). Thus, we turned to the HADDOCK2.4 web server^41^ to generate structures of the complex. The active residues of NpF2164g6 were set to be all those affected by BNp-Red-1.0 addition, and the active residues of BNp-Red-1.0 were set to be those mutated during phage display.

When docking proteins, HADDOCK2.4 generates multiple structures, then groups those structures into clusters based on structural similarity. To validate the docking results, we simulated the top-ranked structures of the top two clusters for 1 µs in the FF14SB force field.^34^ While neither of these structures was stable (Fig. S17, S18), one equilibrated to a structure that was then maintained for the remainder of the simulation. A representative frame of this structure was then selected (see MD methods) and simulated in triplicate for 1 µs. The BNp-Red-1.0 binding orientation was maintained throughout the full simulation in all replicates, in contrast to the initially docked structures (Fig. S17, S18). This structure of the BNp-Red-1.0:NpF2164g6 complex is shown in Figure 3c. A triad of hydrophobic residues (Phe36, Tyr40, and Tyr44) in BNp-Red-1.0 are observed to pack against a hydrophobic portion of NpF2164g6 that includes Trp56, Ile93 and Phe97. In addition, a persistent hydrogen bond is observed between residue Arg44 of BNp-Red-1.0 and Glu59 of NpF2164g6 (Fig. 3c, 96.1± 0.6% occupancy). In all BNp-Red-1.0 derivatives that display photoswitchable NpF2164g6 binding (Fig. S2) Arg44 is maintained, as is the hydrophobic triad. The PISA server (www.ebi.ac.uk/pdbe/pisa/)^54^ calculates a buried surface area of 440 Å^2^ for the interface with residues Phe36 and Tyr40,43 being major contributors to this. The Prodigy webserver,^55^ which calculates the number of interfacial contacts as well as the properties of the non-interacting surfaces,^56^ identifies 30 distinct residue-residue contacts and predicts a binding affinity of −7.7 kcal.mol^−1^ and a dissociation constant at 25 °C of 2.6 µM similar to that measured experimentally (Fig. S19).

### Conformational switching and comparison with phytochrome structures

Figure 4 compares the docked structure of BNp-Red-1.0 and NpF2164g6 with the structures of the Pr and Pfr photostates of the PAS-GAF-PHY domains from *D. radiodurans*, and with the PIF6 bound structure of PhyB.^19, 20, 57^ In all these cases, conformational change is initiated by isomerization of the 15-16 bond in the bilin chromophore. Rearrangements of the adjacent tongue segment of the PHY domain are vital for signal propagation.^58^ The tongue of the PHY domain adopts a β-hairpin conformation in the dark state crystal structure of DrBphP (Fig. 4a)^57^ but solution NMR measurements indicate this region is conformationally heterogeneous.^59^ The tongue adopts a helical conformation in the light (Pfr) state (PDB code: 4O01) (Fig. 4b). The interaction of the plant phytochrome with its binding partner PIF6 in the Pfr state also involves the conversion of the PHY tongue into a helical conformation (Fig. 4c).^19, 20^ In addition, the ∼100 residue PhyB N-terminal domain changes from a disordered state into a set of three helices (olive green in Fig 4c), that interacts with the helical tongue and also with PIF6 (shown in cyan).

**Figure 4.**
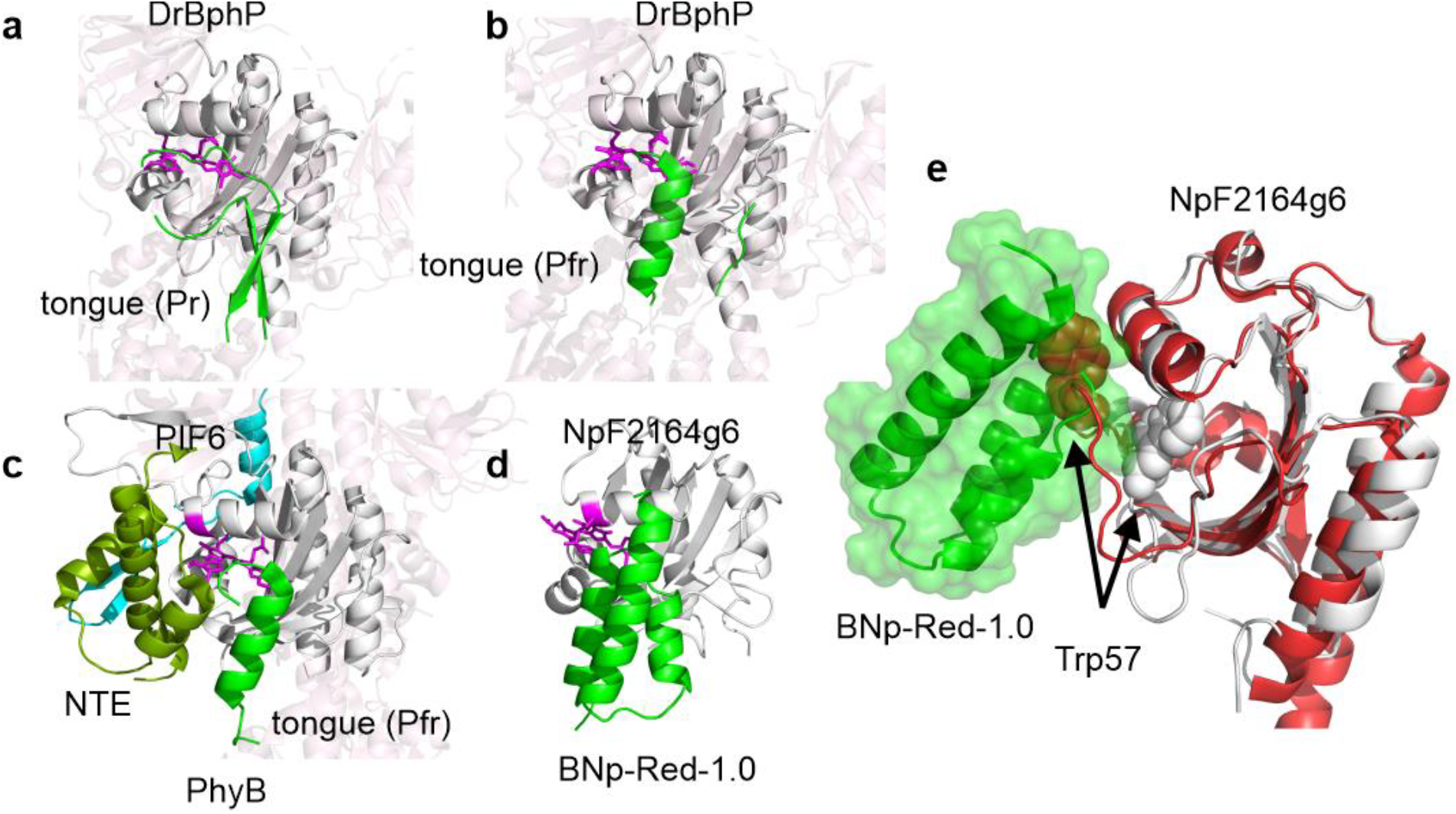
Structures of (a) the dark (Pr) state of bacteriophytochrome DrBph (4O0P), (b) the light state (Pfr) of DrBph (4O01), (c) the Pfr state of the plant phytochrome PhyB bound to PIF6 (cyan) (9JLB), (d) the modeled structure of BNp-Red-1.0 bound to NpF2164g6. The GAF domain in each structure is aligned and shown as a white cartoon with the chromophore in magenta. Other domains in the phytochrome structures are shown as transparent cartoon models. The tongue domain of the phytochromes and the binder BNp-Red-1.0 are shown in green. The N-terminal domain of PhyB that folds upon PIF6 binding to the Pfr state is shown in olive green. ^19, 20^. (e) The BNp-Red-1.0: NpF2164g6 dark state complex. The dark state of NpF2164g6 is shown in white and BNp-Red-1.0 is shown in green. The light state (Pg) structure of the homologous protein NpR6012g4 is shown in red (PDB code: 6BHO). Trp57 is shown in spacefill. If Trp57 moves upon irradiation of NpF2164g6 in the same manner that it does upon irradiation of NpR6012g4 then it would clash with BNp-Red-1.0.

In contrast to these substantial conformational changes in phytochromes, light triggered changes in CBCR domains are more subtle. A detailed analysis of the conformational changes of NpR6012g4 in solution using NMR methods revealed that bilin isomerization causes a shift of residues comprising the extended loop near the chromophore (residues 644 to 655 in NpR6012g4). In the dark state, the Trp655 indole ring (analogous to Trp57 in the NpF2164g6 construct) forms a π-stacking interaction with the bilin D-ring. In the light state, the indole ring moves to become exposed to bulk solvent and close to the C3 side chain on the A-ring (Fig. 4e). A similar rearrangement of this loop and the corresponding Trp residue (Trp496) is seen when comparing the X-ray crystal structures of another CBCR GAF domain Slr1393g3 in the dark state (PDB code: 5DFX) and the light state (PDB code: 5M82).

Our data indicate that the binder BNp-Red-1.0 interacts with a surface of NpF2164g6 that is analogous to the surface in the GAF domain of phytochromes that interacts with the PHY tongue (Fig. 4d). A structural alignment of the GAF domains of phytochromes with that of NpF2164g6 places the helical form of the tongue at a position between helices 2 and 3 (the phage displayed binding interface) of the docked BNp-Red-1.0 binder (Fig. 4). Whereas in the phytochrome structure the tongue undergoes helix/sheet conformational changes, with NpF2164g6 photoswitching directly triggers dissociation/association of BNp-Red-1.0. NMR assignments of the light state structure of NpF2164g6 indicate that the conformational change that occurs is analogous to that seen with NpR6012g4 (Fig. S11). If the loop rearrangement and movement of the Trp residue from a buried to solvent exposed position occurs in NpF2164g6 as is seen in NpR6012g4 and Slr1993g3, the Trp would directly clash with BNp-Red-1.0 (Fig. 4e). Such a conformational change would explain why BNp-Red-1.0 dissociates from NpF2164g6 upon red light absorption and isomerization of the bilin.

Additionally, the Glu59 residue in NpF2164g6 has been reported to be important for tuning the thermal reversion kinetics of red/green CBCR GAF domains.^33^ Specifically, when the side chain of Glu59 is stabilized via H-bonding, enhanced thermal relaxation is observed. The interaction of Arg44 of BNp-Red-1.0 with Glu59 observed in MD simulations (Fig. 3c) suggests that the enhancement of the thermal reversion rate of NpF2164g6 by BNp-Red-1.0 binding may occur by a similar mechanism.

### Summary & Outlook

Here we report a red-light-only optogenetic tool, UNICYCL, based on the 1:1 interaction between the CBCR GAF domain NpF2164g6 and a dark-state selective binder. These components are much smaller than current red-light optogenetic tools and structural characterization of the complex identifies surfaces on both the binder and the photoreceptor that may be targeted for further manipulation of binding affinity and/or binding kinetics. The interface defined here may be further engineered to develop orthogonal pairs of red-light only switches/binders as was done with Magnets^60^ after the Vivid homodimer structure was determined.^61^ Finally, its simple stoichiometry, small size, and structural characterization should facilitate deployment of the UNICYCL tool to enable red-light caging of diverse targets by analogy with the blue light tool Z-lock^62^ and the UV/cyan tool Dronpa.^63^

## Author Contributions

JJ and GNTL performed phage display. JJ performed SEC. PMMR, JJ, and GNTL performed UV-Vis characterization. PMMR performed NMR spectroscopy and molecular dynamics simulations. KT performed the SEAP assay. GNTL performed the colocalization assay. PMMR, GNTL, JJ, KT, MU, MZ, and GAW analyzed data. PMMR, GNTL, and GAW wrote the paper

## References

(1) Goglia, A. G.; Toettcher, J. E. A bright future: optogenetics to dissect the spatiotemporal control of cell behavior. Curr Opin Chem Biol 2019, 48, 106–113. DOI: 10.1016/j.cbpa.2018.11.010 From NLM Medline.

(2) Kolar, K.; Knobloch, C.; Stork, H.; Znidaric, M.; Weber, W. OptoBase: A Web Platform for Molecular Optogenetics. ACS Synth Biol 2018, 7 (7), 1825–1828. DOI: 10.1021/acssynbio.8b00120 From NLM Medline.

(3) Stolik, S.; Delgado, J. A.; Perez, A.; Anasagasti, L. Measurement of the penetration depths of red and near infrared light in human “ex vivo” tissues. J Photochem Photobiol B 2000, 57 (2-3), 90–93. DOI: 10.1016/s1011-1344(00)00082-8 From NLM Medline.

(4) Tang, K.; Beyer, H. M.; Zurbriggen, M. D.; Gartner, W. The Red Edge: Bilin-Binding Photoreceptors as Optogenetic Tools and Fluorescence Reporters. Chem Rev 2021, 121 (24), 14906–14956. DOI: 10.1021/acs.chemrev.1c00194 From NLM Medline.

(5) Redchuk, T. A.; Omelina, E. S.; Chernov, K. G.; Verkhusha, V. V. Near-infrared optogenetic pair for protein regulation and spectral multiplexing. Nat Chem Biol 2017, 13 (6), 633–639. DOI: 10.1038/nchembio.2343 From NLM Medline.

(6) Golonka, D.; Fischbach, P.; Jena, S. G.; Kleeberg, J. R. W.; Essen, L. O.; Toettcher, J. E.; Zurbriggen, M. D.; Moglich, A. Deconstructing and repurposing the light-regulated interplay between Arabidopsis phytochromes and interacting factors. Commun Biol 2019, 2, 448. DOI: 10.1038/s42003-019-0687-9 From NLM PubMed-not-MEDLINE. Machens, F.; Ran, G.; Ruehmkorff, C.; Meyer Auf der Heyde, J.; Mueller-Roeber, B.; Hochrein, L. PhiReX 2.0: A Programmable and Red Light-Regulated CRISPR-dCas9 System for the Activation of Endogenous Genes in Saccharomyces cerevisiae. ACS Synth Biol 2023, 12 (4), 1046–1057. DOI: 10.1021/acssynbio.2c00517 From NLM Medline. Oda, S.; Sato-Ebine, E.; Nakamura, A.; Kimura, K. D.; Aoki, K. Optical Control of Cell Signaling with Red/Far-Red Light-Responsive Optogenetic Tools in Caenorhabditis elegans. ACS Synth Biol 2023, 12 (3), 700–708. DOI: 10.1021/acssynbio.2c00461 From NLM Medline.

(7) Huang, Z.; Li, Z.; Zhang, X.; Kang, S.; Dong, R.; Sun, L.; Fu, X.; Vaisar, D.; Watanabe, K.; Gu, L. Creating Red Light-Switchable Protein Dimerization Systems as Genetically Encoded Actuators with High Specificity. ACS Synth Biol 2020, 9 (12), 3322–3333. DOI: 10.1021/acssynbio.0c00397 From NLM Medline. Zhou, P.; Jia, Y.; Zhang, T.; Abudukeremu, A.; He, X.; Zhang, X.; Liu, C.; Li, W.; Li, Z.; Sun, L.; et al. Red Light-Activated Reversible Inhibition of Protein Functions by Assembled Trap. ACS Synth Biol 2025, 14 (5), 1437–1450. DOI: 10.1021/acssynbio.4c00585 From NLM Medline.

(8) Kuwasaki, Y.; Suzuki, K.; Yu, G.; Yamamoto, S.; Otabe, T.; Kakihara, Y.; Nishiwaki, M.; Miyake, K.; Fushimi, K.; Bekdash, R.; et al. A red light-responsive photoswitch for deep tissue optogenetics. Nat Biotechnol 2022, 40 (11), 1672–1679. DOI: 10.1038/s41587-022-01351-w From NLM Medline.

(9) Zhou, Y.; Kong, D.; Wang, X.; Yu, G.; Wu, X.; Guan, N.; Weber, W.; Ye, H. A small and highly sensitive red/far-red optogenetic switch for applications in mammals. Nat Biotechnol 2022, 40 (2), 262–272. DOI: 10.1038/s41587-021-01036-w From NLM Medline.

(10) Qiao, L.; Niu, L.; Wang, M.; Wang, Z.; Kong, D.; Yu, G.; Ye, H. A sensitive red/far-red photoswitch for controllable gene therapy in mouse models of metabolic diseases. Nat Commun 2024, 15 (1), 10310. DOI: 10.1038/s41467-024-54781-2 From NLM Medline.

(11) Flores-Ibarra, A.; Maia, R. N. A.; Olasz, B.; Church, J. R.; Gotthard, G.; Schapiro, I.; Heberle, J.; Nogly, P. Light-Oxygen-Voltage (LOV)-sensing Domains: Activation Mechanism and Optogenetic Stimulation. J Mol Biol 2024, 436 (5), 168356. DOI: 10.1016/j.jmb.2023.168356 From NLM Medline.

(12) Harper, S. M.; Neil, L. C.; Gardner, K. H. Structural basis of a phototropin light switch. Science 2003, 301 (5639), 1541–1544. DOI: 10.1126/science.1086810 From NLM Medline. Moglich, A.; Moffat, K. Engineered photoreceptors as novel optogenetic tools. Photochem Photobiol Sci 2010, 9 (10), 1286–1300. DOI: 10.1039/c0pp00167h From NLM Medline.

(13) Motta-Mena, L. B.; Reade, A.; Mallory, M. J.; Glantz, S.; Weiner, O. D.; Lynch, K. W.; Gardner, K. H. An optogenetic gene expression system with rapid activation and deactivation kinetics. Nat Chem Biol 2014, 10 (3), 196–202. DOI: 10.1038/nchembio.1430 From NLM Medline.

(14) Richter, F.; Fonfara, I.; Bouazza, B.; Schumacher, C. H.; Bratovic, M.; Charpentier, E.; Moglich, A. Engineering of temperature- and light-switchable Cas9 variants. Nucleic Acids Res 2016, 44 (20), 10003–10014. DOI: 10.1093/nar/gkw930 From NLM Medline.

(15) Li, X. L.; Tei, R.; Uematsu, M.; Baskin, J. M. Ultralow Background Membrane Editors for Spatiotemporal Control of Phosphatidic Acid Metabolism and Signaling. ACS Cent Sci 2024, 10 (3), 543–554. DOI: 10.1021/acscentsci.3c01105 From NLM PubMed-not-MEDLINE.

(16) Yumerefendi, H.; Dickinson, D. J.; Wang, H.; Zimmerman, S. P.; Bear, J. E.; Goldstein, B.; Hahn, K.; Kuhlman, B. Control of Protein Activity and Cell Fate Specification via Light-Mediated Nuclear Translocation. PLoS One 2015, 10 (6), e0128443. DOI: 10.1371/journal.pone.0128443 From NLM Medline. Niopek, D.; Wehler, P.; Roensch, J.; Eils, R.; Di Ventura, B. Optogenetic control of nuclear protein export. Nat Commun 2016, 7, 10624. DOI: 10.1038/ncomms10624 From NLM Medline.

(17) Strickland, D.; Yao, X.; Gawlak, G.; Rosen, M. K.; Gardner, K. H.; Sosnick, T. R. Rationally improving LOV domain-based photoswitches. Nat Methods 2010, 7 (8), 623–626. DOI: 10.1038/nmeth.1473 From NLM Medline. Pudasaini, A.; El-Arab, K. K.; Zoltowski, B. D. LOV-based optogenetic devices: light-driven modules to impart photoregulated control of cellular signaling. Front Mol Biosci 2015, 2, 18. DOI: 10.3389/fmolb.2015.00018 From NLM PubMed-not-MEDLINE. Niu, J.; Ben Johny, M.; Dick, I. E.; Inoue, T. Following Optogenetic Dimerizers and Quantitative Prospects. Biophys J 2016, 111 (6), 1132–1140. DOI: 10.1016/j.bpj.2016.07.040 From NLM Medline.

(18) Kaeser, G.; Krauss, N.; Roughan, C.; Sauthof, L.; Scheerer, P.; Lamparter, T. Phytochrome-Interacting Proteins. Biomolecules 2023, 14 (1). DOI: 10.3390/biom14010009 From NLM Medline. Multamaki, E.; Nanekar, R.; Morozov, D.; Lievonen, T.; Golonka, D.; Wahlgren, W. Y.; Stucki-Buchli, B.; Rossi, J.; Hytonen, V. P.; Westenhoff, S.; et al. Comparative analysis of two paradigm bacteriophytochromes reveals opposite functionalities in two-component signaling. Nat Commun 2021, 12 (1), 4394. DOI: 10.1038/s41467-021-24676-7 From NLM Medline.

(19) Wang, Z.; Wang, W.; Zhao, D.; Song, Y.; Lin, X.; Shen, M.; Chi, C.; Xu, B.; Zhao, J.; Deng, X. W.; et al. Light-induced remodeling of phytochrome B enables signal transduction by phytochrome-interacting factor. Cell 2024, 187 (22), 6235–6250 e6219. DOI: 10.1016/j.cell.2024.09.005 From NLM Medline.

(20) Jia, H.; Guan, Z.; Ding, J.; Wang, X.; Tian, D.; Zhu, Y.; Zhang, D.; Liu, Z.; Ma, L.; Yin, P. Structural insight into PIF6-mediated red light signal transduction of plant phytochrome B. Cell Discov 2025, 11 (1), 51. DOI: 10.1038/s41421-025-00802-3 From NLM PubMed-not-MEDLINE.

(21) Smith, R. W.; Helwig, B.; Westphal, A. H.; Pel, E.; Borst, J. W.; Fleck, C. Interactions Between phyB and PIF Proteins Alter Thermal Reversion Reactions in vitro. Photochem Photobiol 2017, 93 (6), 1525–1531. DOI: 10.1111/php.12793 From NLM Medline.

(22) Jang, J.; Tang, K.; Youn, J.; McDonald, S.; Beyer, H. M.; Zurbriggen, M. D.; Uppalapati, M.; Woolley, G. A. Engineering of bidirectional, cyanobacteriochrome-based light-inducible dimers (BICYCL)s. Nat Methods 2023, 20 (3), 432–441. DOI: 10.1038/s41592-023-01764-8 From NLM Medline.

(23) Le, G. N. T.; Jang, J.; Uppalapati, M.; Woolley, G. A. Optimized Phage Display-Based Selection for the Development of Heterodimerizing Optogenetic Tools. ACS Synth Biol 2025, 14 (6), 2400–2404. DOI: 10.1021/acssynbio.5c00167 From NLM Medline.

(24) Reis, J. M.; Xu, X.; McDonald, S.; Woloschuk, R. M.; Jaikaran, A. S. I.; Vizeacoumar, F. S.; Woolley, G. A.; Uppalapati, M. Discovering Selective Binders for Photoswitchable Proteins Using Phage Display. ACS Synth Biol 2018, 7 (10), 2355–2364. DOI: 10.1021/acssynbio.8b00123 From NLM Medline.

(25) Fushimi, K.; Enomoto, G.; Ikeuchi, M.; Narikawa, R. Distinctive Properties of Dark Reversion Kinetics between Two Red/Green-Type Cyanobacteriochromes and their Application in the Photoregulation of cAMP Synthesis. Photochem Photobiol 2017, 93 (3), 681–691. DOI: 10.1111/php.12732 From NLM Medline.

(26) Rockwell, N. C.; Martin, S. S.; Lagarias, J. C. Red/green cyanobacteriochromes: sensors of color and power. Biochemistry 2012, 51 (48), 9667–9677. DOI: 10.1021/bi3013565 From NLM Medline.

(27) Gambetta, G. A.; Lagarias, J. C. Genetic engineering of phytochrome biosynthesis in bacteria. Proc Natl Acad Sci U S A 2001, 98 (19), 10566–10571. DOI: 10.1073/pnas.191375198 From NLM Medline.

(28) Fairhead, M.; Howarth, M. Site-specific biotinylation of purified proteins using BirA. Methods Mol Biol 2015, 1266, 171–184. DOI: 10.1007/978-1-4939-2272-7_12 From NLM Medline.

(29) Horner, M.; Gerhardt, K.; Salavei, P.; Hoess, P.; Harrer, D.; Kaiser, J.; Tabor, J. J.; Weber, W. Production of Phytochromes by High-Cell-Density E. coli Fermentation. ACS Synth Biol 2019, 8 (10), 2442–2450. DOI: 10.1021/acssynbio.9b00267 From NLM Medline.

(30) Glazer, A. N.; Fang, S. Chromophore content of blue-green algal phycobiliproteins. J Biol Chem 1973, 248 (2), 659–662. From NLM Medline.

(31) Clubb, R. T.; Thanabal, V.; Wagner, G. A constant-time three-dimensional triple-resonance pulse scheme to correlate intraresidue 1HN, 15N, and 13C′ chemical shifts in 15N-13C-labelled proteins. J. Magn. Reson. 1992, 97 (1), 213–217.

(32) Reed, P. M. M.; Jang, J.; Woloschuk, R. M.; Reis, J.; Hille, J. I. C.; Uppalapati, M.; Woolley, G. A. Effects of binding partners on thermal reversion rates of photoswitchable molecules. Proc Natl Acad Sci U S A 2025, 122 (10), e2414748122. DOI: 10.1073/pnas.2414748122 From NLM Medline.

(33) Jang, J.; Reed, P. M. M.; Rauscher, S.; Woolley, G. A. Point (S-to-G) Mutations in the W(S/G)GE Motif in Red/Green Cyanobacteriochrome GAF Domains Enhance Thermal Reversion Rates. Biochemistry 2022, 61 (14), 1444–1455. DOI: 10.1021/acs.biochem.2c00060 From NLM Medline.

(34) Maier, J. A.; Martinez, C.; Kasavajhala, K.; Wickstrom, L.; Hauser, K. E.; Simmerling, C. ff14SB: Improving the Accuracy of Protein Side Chain and Backbone Parameters from ff99SB. J Chem Theory Comput 2015, 11 (8), 3696–3713. DOI: 10.1021/acs.jctc.5b00255 From NLM Medline.

(35) Wang, J.; Wolf, R. M.; Caldwell, J. W.; Kollman, P. A.; Case, D. A. Development and testing of a general amber force field. J Comput Chem 2004, 25 (9), 1157–1174. DOI: 10.1002/jcc.20035 From NLM Medline.

(36) Rao, A. G.; Wiebeler, C.; Sen, S.; Cerutti, D. S.; Schapiro, I. Histidine protonation controls structural heterogeneity in the cyanobacteriochrome AnPixJg2. Phys Chem Chem Phys 2021, 23 (12), 7359–7367. DOI: 10.1039/d0cp05314g From NLM Medline.

(37) AMBER 2018; University of California, San Francisco: 2018. (accessed.

(38) Kim, G.; Lee, S.; Levy Karin, E.; Kim, H.; Moriwaki, Y.; Ovchinnikov, S.; Steinegger, M.; Mirdita, M. Easy and accurate protein structure prediction using ColabFold. Nat Protoc 2025, 20 (3), 620–642. DOI: 10.1038/s41596-024-01060-5 From NLM Medline.

(39) Narikawa, R.; Ishizuka, T.; Muraki, N.; Shiba, T.; Kurisu, G.; Ikeuchi, M. Structures of cyanobacteriochromes from phototaxis regulators AnPixJ and TePixJ reveal general and specific photoconversion mechanism. Proc Natl Acad Sci U S A 2013, 110 (3), 918–923. DOI: 10.1073/pnas.1212098110 From NLM Medline.

(40) Roe, D. R.; Cheatham, T. E., 3rd. PTRAJ and CPPTRAJ: Software for Processing and Analysis of Molecular Dynamics Trajectory Data. J Chem Theory Comput 2013, 9 (7), 3084–3095. DOI: 10.1021/ct400341p From NLM PubMed-not-MEDLINE.

(41) Honorato, R. V.; Koukos, P. I.; Jimenez-Garcia, B.; Tsaregorodtsev, A.; Verlato, M.; Giachetti, A.; Rosato, A.; Bonvin, A. Structural Biology in the Clouds: The WeNMR-EOSC Ecosystem. Front Mol Biosci 2021, 8, 729513. DOI: 10.3389/fmolb.2021.729513 From NLM PubMed-not-MEDLINE. Honorato, R. V.; Trellet, M. E.; Jimenez-Garcia, B.; Schaarschmidt, J. J.; Giulini, M.; Reys, V.; Koukos, P. I.; Rodrigues, J.; Karaca, E.; van Zundert, G. C. P.; et al. The HADDOCK2.4 web server for integrative modeling of biomolecular complexes. Nat Protoc 2024, 19 (11), 3219–3241. DOI: 10.1038/s41596-024-01011-0 From NLM Medline.

(42) Mates, L.; Chuah, M. K.; Belay, E.; Jerchow, B.; Manoj, N.; Acosta-Sanchez, A.; Grzela, D. P.; Schmitt, A.; Becker, K.; Matrai, J.; et al. Molecular evolution of a novel hyperactive Sleeping Beauty transposase enables robust stable gene transfer in vertebrates. Nat Genet 2009, 41 (6), 753–761. DOI: 10.1038/ng.343 From NLM Medline.

(43) Kowarz, E.; Loscher, D.; Marschalek, R. Optimized Sleeping Beauty transposons rapidly generate stable transgenic cell lines. Biotechnol J 2015, 10 (4), 647–653. DOI: 10.1002/biot.201400821 From NLM Medline.

(44) Beyer, H. M.; Kumar, S.; Nieke, M.; Diehl, C. M. C.; Tang, K.; Shumka, S.; Koh, C. S.; Fleck, C.; Davies, J. A.; Khammash, M.; et al. Genetically-stable engineered optogenetic gene switches modulate spatial cell morphogenesis in two- and three-dimensional tissue cultures. Nat Commun 2024, 15 (1), 10470. DOI: 10.1038/s41467-024-54350-7 From NLM Medline.

(45) Chang, C. W.; Gottlieb, S. M.; Kim, P. W.; Rockwell, N. C.; Lagarias, J. C.; Larsen, D. S. Reactive ground-state pathways are not ubiquitous in red/green cyanobacteriochromes. J Phys Chem B 2013, 117 (38), 11229–11238. DOI: 10.1021/jp402112u From NLM Medline.

(46) Heyne, K.; Herbst, J.; Stehlik, D.; Esteban, B.; Lamparter, T.; Hughes, J.; Diller, R. Ultrafast dynamics of phytochrome from the cyanobacterium synechocystis, reconstituted with phycocyanobilin and phycoerythrobilin. Biophys J 2002, 82 (2), 1004–1016. DOI: 10.1016/S0006-3495(02)75460-X From NLM Medline. Muller, M. G.; Lindner, I.; Martin, I.; Gartner, W.; Holzwarth, A. R. Femtosecond kinetics of photoconversion of the higher plant photoreceptor phytochrome carrying native and modified chromophores. Biophys J 2008, 94 (11), 4370–4382. DOI: 10.1529/biophysj.106.091652 From NLM Medline. Consiglieri, E.; Gutt, A.; Gartner, W.; Schubert, L.; Viappiani, C.; Abbruzzetti, S.; Losi, A. Dynamics and efficiency of photoswitching in biliverdin-binding phytochromes. Photochem Photobiol Sci 2019, 18 (10), 2484–2496. DOI: 10.1039/c9pp00264b From NLM PubMed-not-MEDLINE.

(47) Guinn, M. T.; Coraci, D.; Guinn, L.; Balazsi, G. Reliably Engineering and Controlling Stable Optogenetic Gene Circuits in Mammalian Cells. J Vis Exp 2021, (173). DOI: 10.3791/62109 From NLM Medline.

(48) Guntas, G.; Hallett, R. A.; Zimmerman, S. P.; Williams, T.; Yumerefendi, H.; Bear, J. E.; Kuhlman, B. Engineering an improved light-induced dimer (iLID) for controlling the localization and activity of signaling proteins. Proc Natl Acad Sci U S A 2015, 112 (1), 112–117. DOI: 10.1073/pnas.1417910112 From NLM Medline.

(49) Lim, S.; Yu, Q.; Gottlieb, S. M.; Chang, C. W.; Rockwell, N. C.; Martin, S. S.; Madsen, D.; Lagarias, J. C.; Larsen, D. S.; Ames, J. B. Correlating structural and photochemical heterogeneity in cyanobacteriochrome NpR6012g4. Proc Natl Acad Sci U S A 2018, 115 (17), 4387–4392. DOI: 10.1073/pnas.1720682115 From NLM Medline. Lim, S.; Yu, Q.; Rockwell, N. C.; Martin, S. S.; Lagarias, J. C.; Ames, J. B. 1H, 13C, and 15N chemical shift assignments of cyanobacteriochrome NpR6012g4 in the green-absorbing photoproduct state. Biomol NMR Assign 2016, 10 (1), 157–161. DOI: 10.1007/s12104-015-9657-4 From NLM Medline.

(50) Mariani, V.; Biasini, M.; Barbato, A.; Schwede, T. lDDT: a local superposition-free score for comparing protein structures and models using distance difference tests. Bioinformatics 2013, 29 (21), 2722–2728. DOI: 10.1093/bioinformatics/btt473 From NLM Medline.

(51) Shen, Y.; Bax, A. Protein backbone and sidechain torsion angles predicted from NMR chemical shifts using artificial neural networks. J Biomol NMR 2013, 56 (3), 227–241. DOI: 10.1007/s10858-013-9741-y From NLM Medline.

(52) Heo, L.; Feig, M. Experimental accuracy in protein structure refinement via molecular dynamics simulations. Proc Natl Acad Sci U S A 2018, 115 (52), 13276–13281. DOI: 10.1073/pnas.1811364115 From NLM Medline.

(53) Abramson, J.; Adler, J.; Dunger, J.; Evans, R.; Green, T.; Pritzel, A.; Ronneberger, O.; Willmore, L.; Ballard, A. J.; Bambrick, J.; et al. Accurate structure prediction of biomolecular interactions with AlphaFold 3. Nature 2024, 630 (8016), 493–500. DOI: 10.1038/s41586-024-07487-w From NLM Medline.

(54) Krissinel, E.; Henrick, K. Inference of macromolecular assemblies from crystalline state. J Mol Biol 2007, 372 (3), 774–797. DOI: 10.1016/j.jmb.2007.05.022 From NLM Medline.

(55) Xue, L. C.; Rodrigues, J. P.; Kastritis, P. L.; Bonvin, A. M.; Vangone, A. PRODIGY: a web server for predicting the binding affinity of protein-protein complexes. Bioinformatics 2016, 32 (23), 3676–3678. DOI: 10.1093/bioinformatics/btw514 From NLM Medline.

(56) Kastritis, P. L.; Rodrigues, J. P.; Folkers, G. E.; Boelens, R.; Bonvin, A. M. Proteins feel more than they see: fine-tuning of binding affinity by properties of the non-interacting surface. J Mol Biol 2014, 426 (14), 2632–2652. DOI: 10.1016/j.jmb.2014.04.017 From NLM Medline.

(57) Takala, H.; Bjorling, A.; Berntsson, O.; Lehtivuori, H.; Niebling, S.; Hoernke, M.; Kosheleva, I.; Henning, R.; Menzel, A.; Ihalainen, J. A.; et al. Signal amplification and transduction in phytochrome photosensors. Nature 2014, 509 (7499), 245–248. DOI: 10.1038/nature13310 From NLM Medline.

(58) Kurttila, M.; Rumfeldt, J.; Takala, H.; Ihalainen, J. A. The interconnecting hairpin extension “arm”: An essential allosteric element of phytochrome activity. Structure 2023, 31 (9), 1100–1108 e1104. DOI: 10.1016/j.str.2023.06.007 From NLM Medline.

(59) Gustavsson, E.; Isaksson, L.; Persson, C.; Mayzel, M.; Brath, U.; Vrhovac, L.; Ihalainen, J. A.; Karlsson, B. G.; Orekhov, V.; Westenhoff, S. Modulation of Structural Heterogeneity Controls Phytochrome Photoswitching. Biophys J 2020, 118 (2), 415–421. DOI: 10.1016/j.bpj.2019.11.025 From NLM Medline.

(60) Kawano, F.; Suzuki, H.; Furuya, A.; Sato, M. Engineered pairs of distinct photoswitches for optogenetic control of cellular proteins. Nat Commun 2015, 6, 6256. DOI: 10.1038/ncomms7256 From NLM Medline.

(61) Vaidya, A. T.; Chen, C. H.; Dunlap, J. C.; Loros, J. J.; Crane, B. R. Structure of a light-activated LOV protein dimer that regulates transcription. Sci Signal 2011, 4 (184), ra50. DOI: 10.1126/scisignal.2001945 From NLM Medline.

(62) Stone, O. J.; Pankow, N.; Liu, B.; Sharma, V. P.; Eddy, R. J.; Wang, H.; Putz, A. T.; Teets, F. D.; Kuhlman, B.; Condeelis, J. S.; et al. Optogenetic control of cofilin and alphaTAT in living cells using Z-lock. Nat Chem Biol 2019, 15 (12), 1183–1190. DOI: 10.1038/s41589-019-0405-4 From NLM Medline.

(63) Zhou, X. X.; Chung, H. K.; Lam, A. J.; Lin, M. Z. Optical control of protein activity by fluorescent protein domains. Science 2012, 338 (6108), 810–814. DOI: 10.1126/science.1226854 From NLM Medline.

