## Supplemental Information for "Red-light-only control of protein-protein interactions using a cyanobacteriochrome (UNICYCL)"

### Supplementary Information:

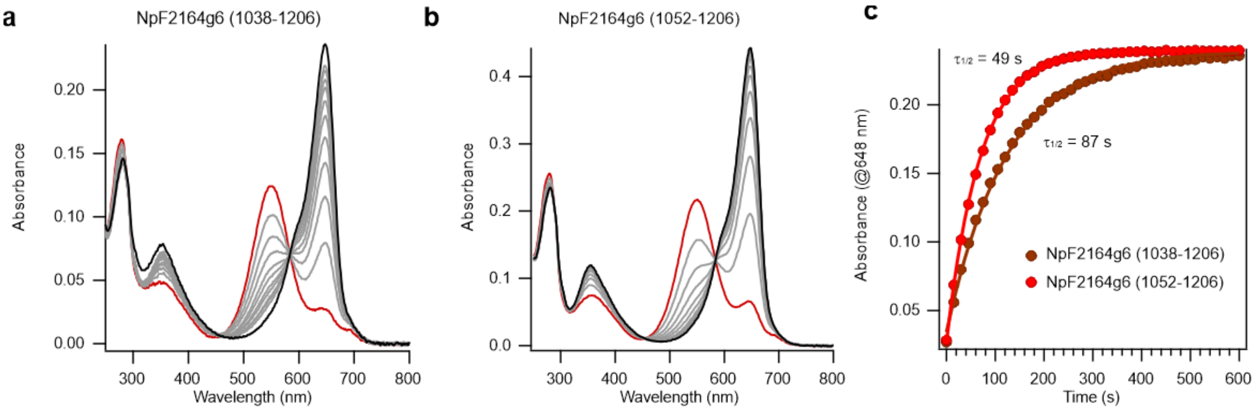


**Figure S1.** Dark reversion of NpF2164g6. (a) The UV-Vis spectrum of dark-adapted NpF2164g6 (1038-1206) (black line), then immediately after red light illumination (660 nm) (red line), then at 30 s intervals during thermal reversion (grey lines). (b) UV-Vis spectrum of truncated dark-adapted NpF2164g6 (1052-1206) (black line), then immediately after red light illumination (red line), then at 30 s intervals during thermal reversion (grey lines). (c) Absorbance at 648 nm monitored over time during thermal reversion for NpF2164g6 (1038-1206) (dark red symbols) and the truncated version NpF2164g6 (1052-1206) (bright red symbols). Single exponential fits are shown with calculated half-lives indicated. The Pg state thermally reverts to the Pr state with a half-life of ~1 minute.

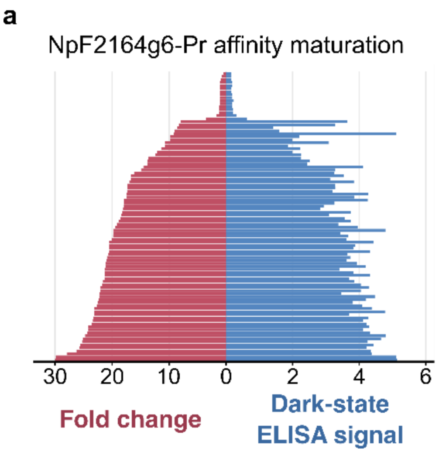


b

| #Position | 25 | 27 | 28 | 31 | 34 | 36 | 37 | 39 | 40 | 43 | 44 | 47 | Fold change | |
|---|---|---|---|---|---|---|---|---|---|---|---|---|---|---|
| A12 | A | A | Y | W | N | W | D | D | Y | Y | R | K | 30.0 | BNp-Red-1.2 |
| A11 | A | A | Y | W | N | W | D | D | Y | Y | R | K | 29.9 | |
| A09 | A | A | Y | L | D | W | D | D | Y | Y | R | K | 27.9 | BNp-Red-1.1 |
| A10 | A | A | Y | L | D | W | D | D | Y | Y | R | K | 26.2 | |
| B01 | A | A | Y | W | N | W | D | D | Y | Y | R | K | 25.7 | |
| C01 | A | A | Y | W | N | Y | D | D | Y | Y | R | K | 22.2 | |
| E1 | S | A | Y | W | N | W | D | D | Y | Y | R | K | 21.3 | |
| G02 | S | A | Y | L | N | W | D | D | Y | Y | R | K | 18.3 | |
| A3 | S | A | Y | V | Q | W | D | D | Y | Y | R | K | 18.1 | |
| F1 | S | A | Y | L | M | W | D | D | Y | Y | R | K | 18.0 | |
| C12 | A | A | Y | W | N | Y | D | D | Y | Y | R | K | 13.7 | |
| B11 | A | A | Y | L | N | I | D | D | Y | Y | R | K | 12.1 | |
| F2 | A | A | Y | L | N | W | D | H | Y | Y | R | K | 8.6 | |

**Figure S2:** (a) Single-clone ELISA results presented as butterfly plot. For each of the 96 clones, the dark-state ELISA signal (right side) is shown alongside its fold change (left side), calculated as the ratio of dark-state to light-state ELISA signal. (b) Amino acid sequences of the randomized residues from representative clones exhibiting a range of fold changes. Numbering is based on PDBcode:1TF0.

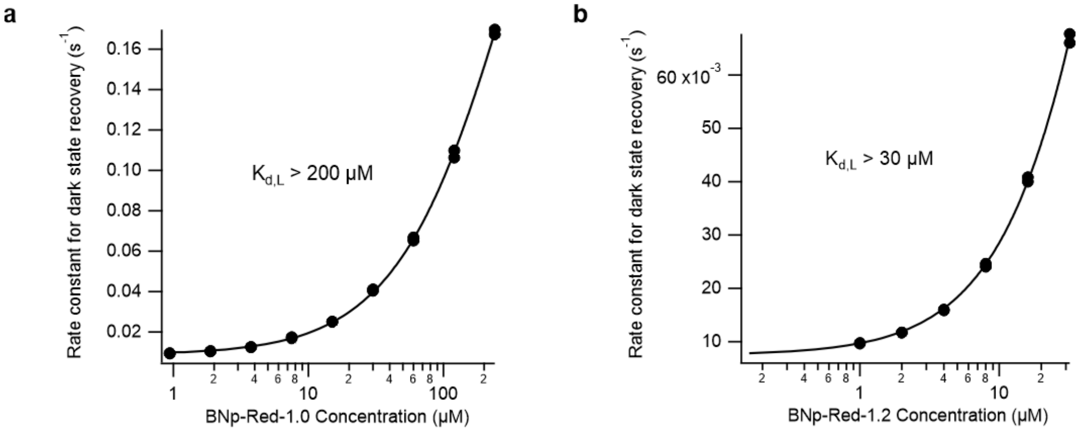


**Figure S3:** The rate constant for thermal reversion observed for NpF2164g6 versus the concentration of (a) BNp-Red-1.0 and (b) BNp-Red-1.2. Complete saturation could not be accomplished, but these data indicate a very weak interaction of the binders with the light state.

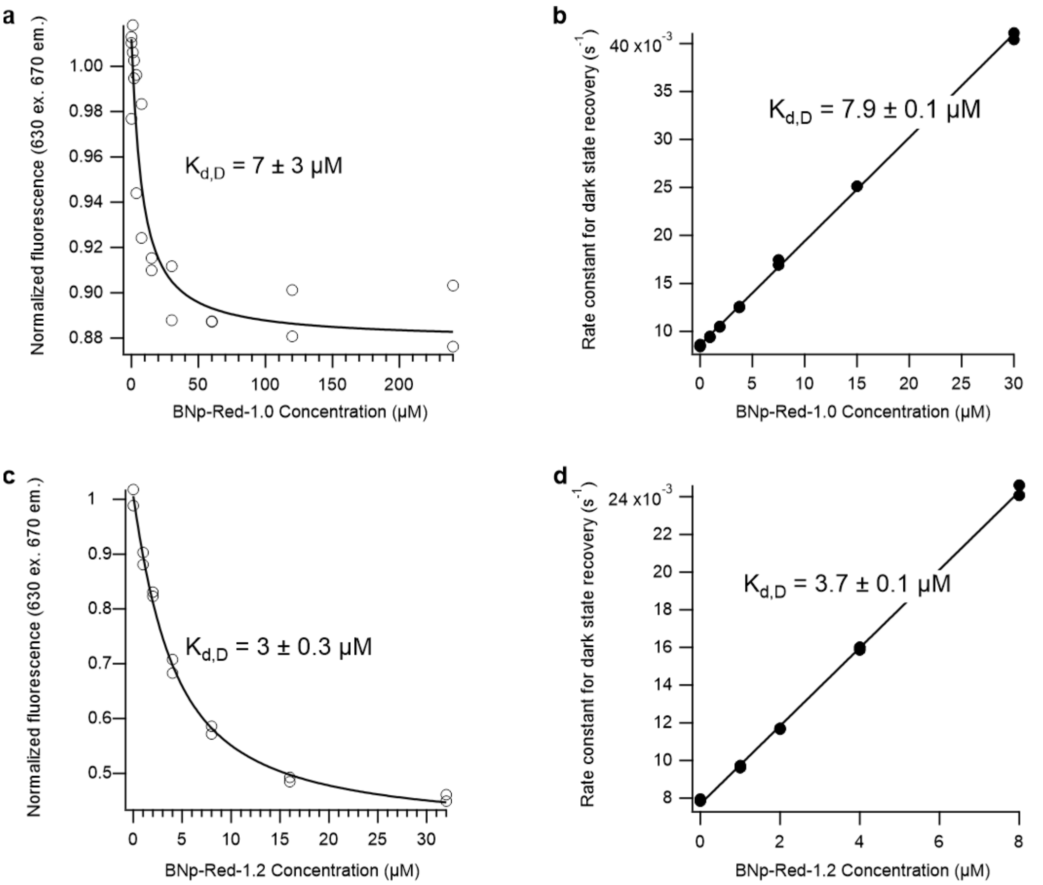


**Figure S4:** Thermal reversion and fluorescence characterization of NpF2164g6. (a,c) Fluorescence of the Pr state of NpF2164g6 versus concentration of the (a) BNp-Red-1.0 binder and (c) the BNp-Red-1.2 binder. The titration yields the $K_{d,D}$. (b,d) Rate constant for thermal reversion of NpF2164g6 versus the concentration of (b) BNp-Red-1.0 and (d) BNp-Red-1.2 for low binder concentrations where the concentration of binder is much smaller than $K_{d,L.}$ These lines can be fit to yield $K_{d,D}$ on the assumption that the interactions have $\phi_{switching}$ values of 1. The close match of the $K_{d,D}$ values from (a-b) and (c-d) support this assumption.

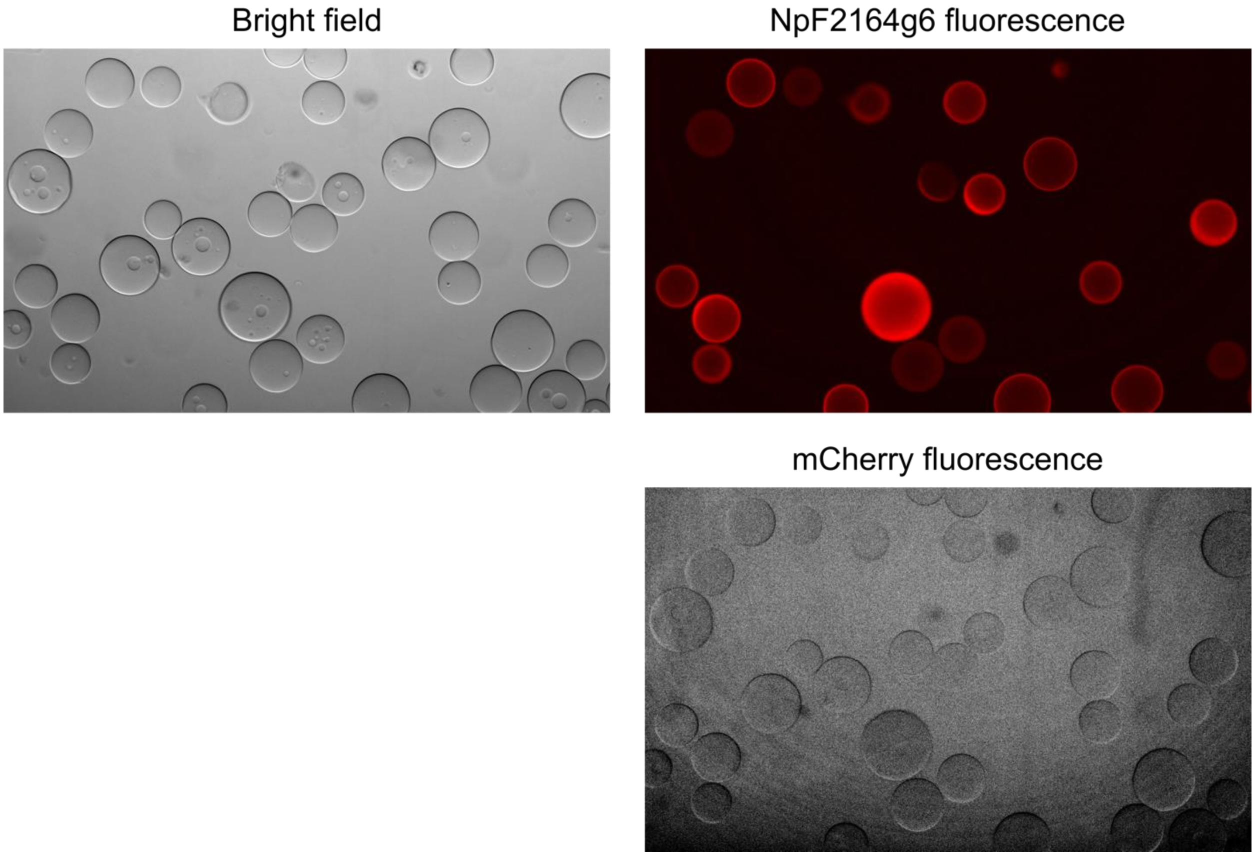


**Figure S5:** A mixture of NpF2164g6-coated and IgG-coated (negative control) agarose beads were resuspended in a solution containing 500 nM mCherry-BNp-Red-1.2. Bright-field image (top left) and NpF2164g6 fluorescence image (top right) of NpF2164g6-coated and IgG-coated beads. mCherry fluorescence image (bottom) in the dark exhibit no localization of mCherry-BNp-Red-1.2 on the surface of NpF2164g6-coated beads.

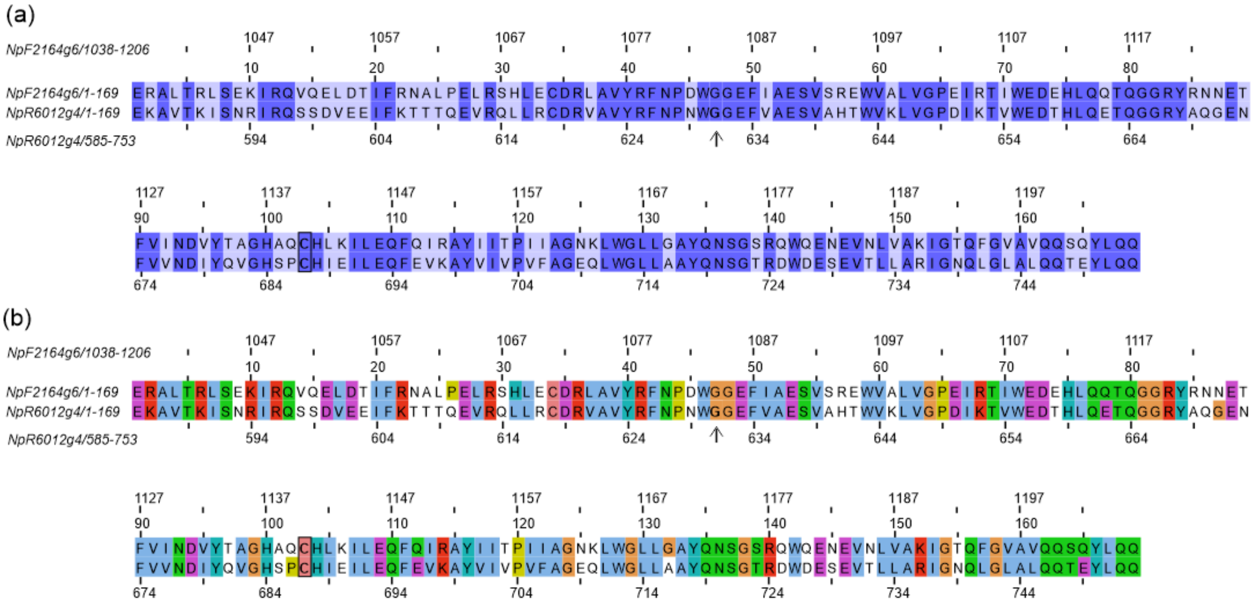


**Figure S6.** Alignment of NpF2164g6 (UniProt: B2J668 · B2J668_NOSP7; NCBI Reference Sequence: WP_012408769.1 (residues 1038-1206) and NpR6012g4 (UniProt: B2IU14 · B2IU14_NOSP7; NCBI Reference Sequence: WP_012412243.1) (residues 585-753) (T631G). Residue 1 in the NpF2164g6 numbering above is residue 1038 in the full protein numbering, while residue 2 in the truncated NpF2164g6 numbering is residue 1052. The site of the T631G mutation is indicated by an arrow and bolded in NpR6012g4. The chromophore-ligating cysteine is also bolded and boxed in both sequences. The proteins are highly homologous, with 59.2% identity (a) and 83.4% similarity (b).

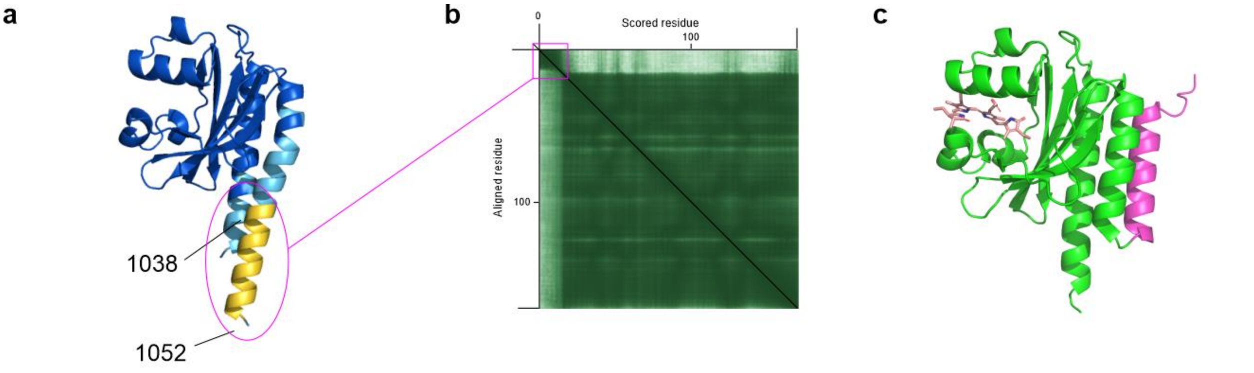


**Figure S7.** (a) Alphafold3 model of NpF2164g6 (1038-1206) coloroued by pLDDT score, where yellow is pLDDT<50 (low confidence). (b) These residues (1038-1051) also show a high predicted aligned error (PAE) vs. the rest of the model (boxed region in magenta). (c). In the crystal structure AnPixJg2 (3W2Z), the region homologous to NpF2164g6 1038-1051 (coloured in magenta) packs back against the N- and C-terminal helices.

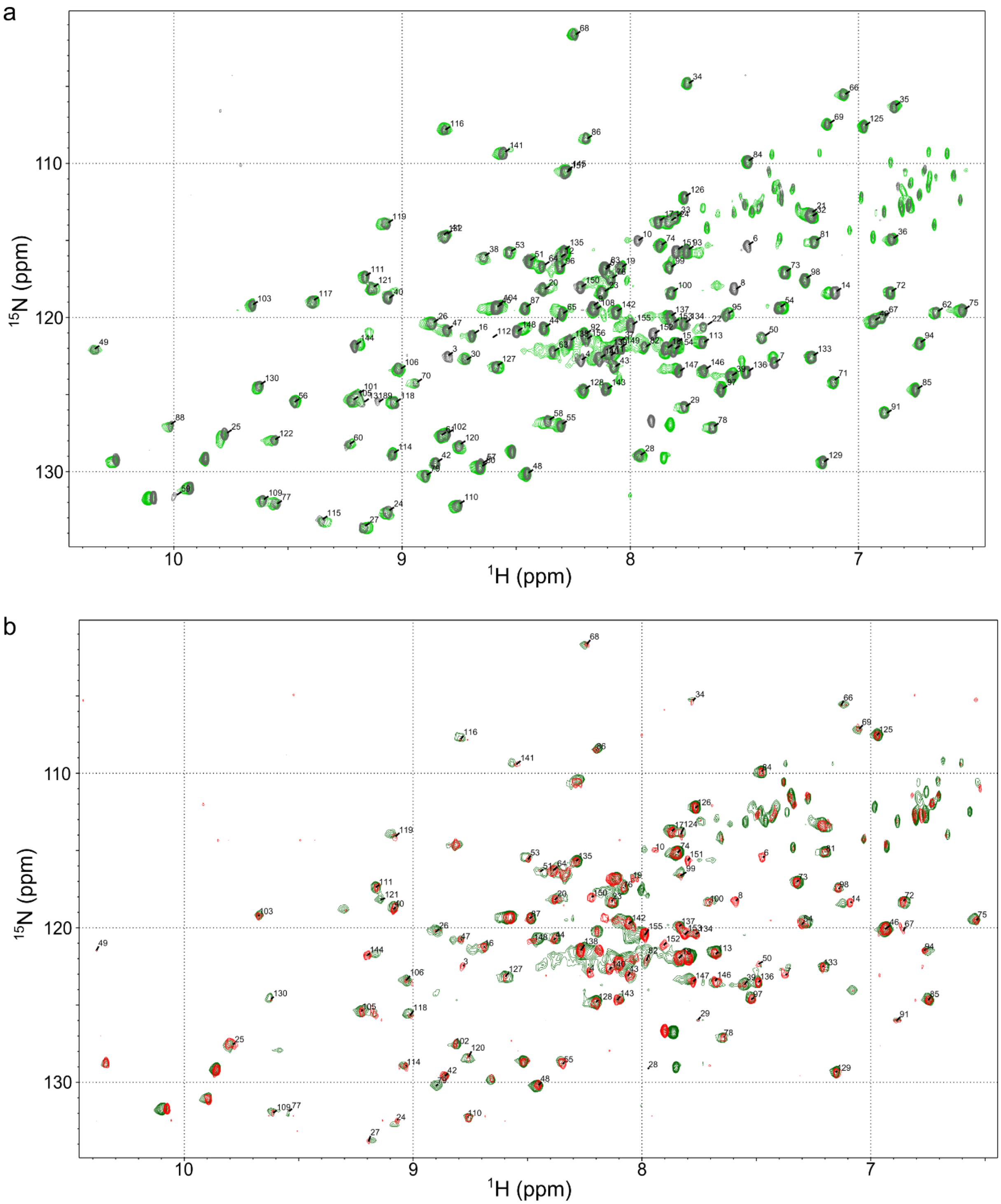


**Figure S8**. NMR spectra comparing dark state NpF2164g6 (1038-1206) to truncated NpF2164g6 (1052-1206). The primary difference is that the longer construct displays several additional peaks in the centre of the spectrum. (a) Truncated (grey) and full length (green) NpF2164g6 alone. (b) Truncated (red) and full length (green) NpF2164g6 in the presence of a molar excess of BNp-Red-1.0.

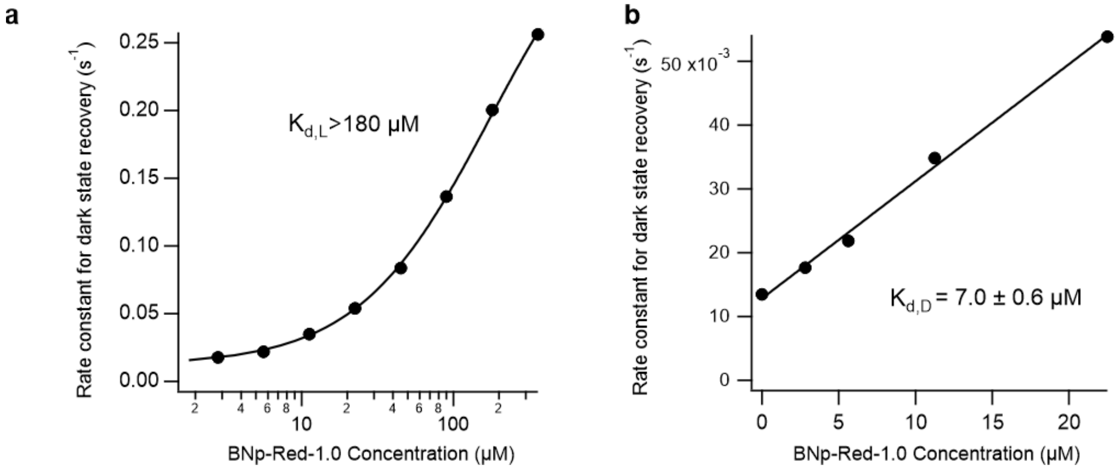


**Figure S9**. Thermal reversion of truncated NpF2164g6 with BNp-Red-1.0. (a) The rate constant of thermal reversion observed for NpF2164g6 versus the concentration of BNp-Red-1.0 (compare to Fig. S3a). (b) Rate constant of thermal reversion of NpF2164g6 versus the concentration of BNp-Red-1.0 for low binder concentrations where the concentration of binder is much smaller than $K_{d,L.}$ This trend can be fit to yield $K_{d,D}$ on the assumption that the interaction has a $\phi_{switching}$ value of 1. Compare to Fig. S4b.

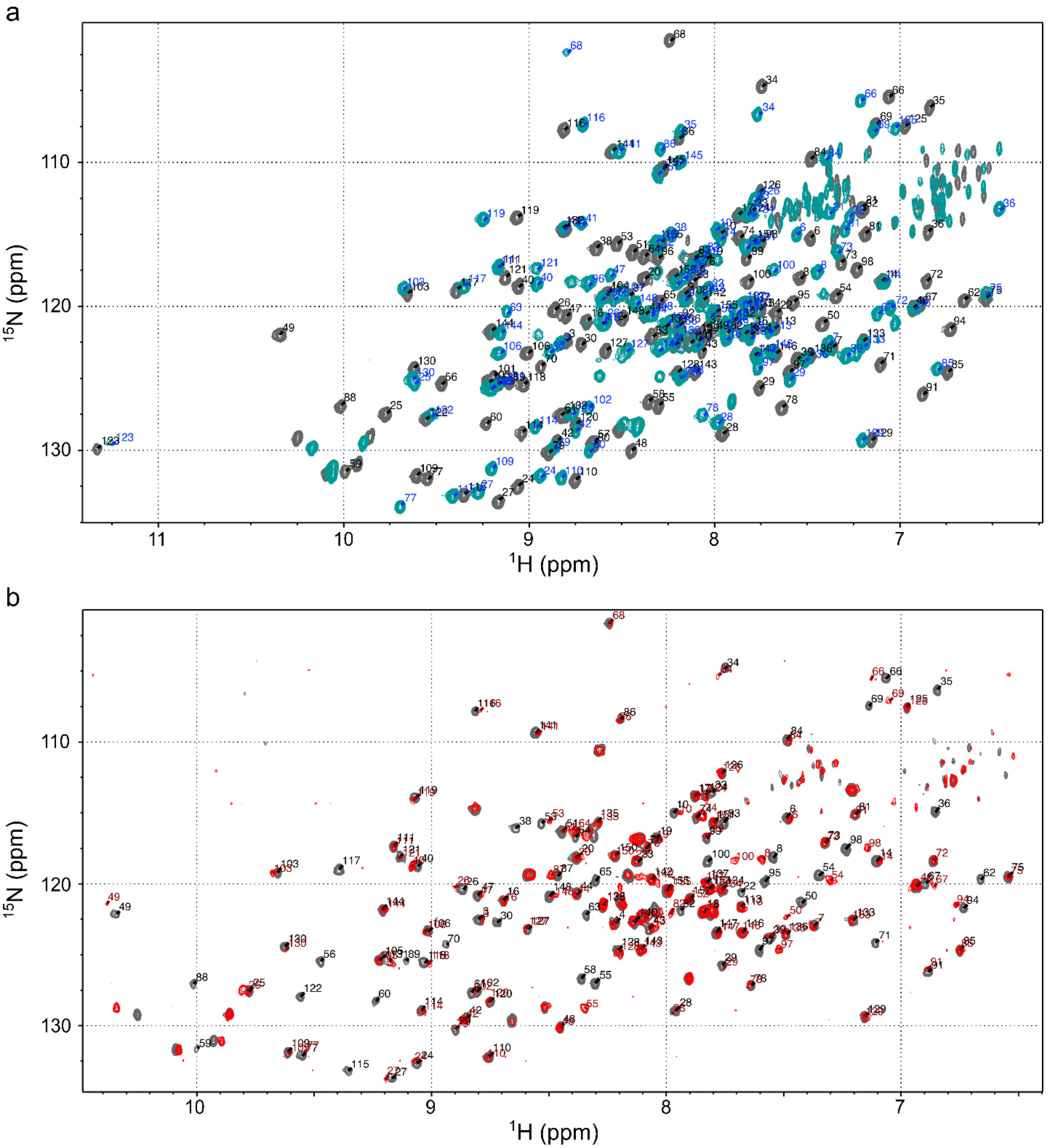


**Figure S10.** (a) NMR spectra of NpF2164g6 (1052-1206) in the dark (grey) and under 660 nm light irradiation (cyan). The assignments are shown in black for the dark state and blue for the light state. (b) NMR spectra of NpF2164g6 (1052-1206) alone in the dark (grey) and in the dark in the presence of a molar excess of BNp-Red-1.0 (red). Assignments were transferred to the latter by inspection and are shown in red.

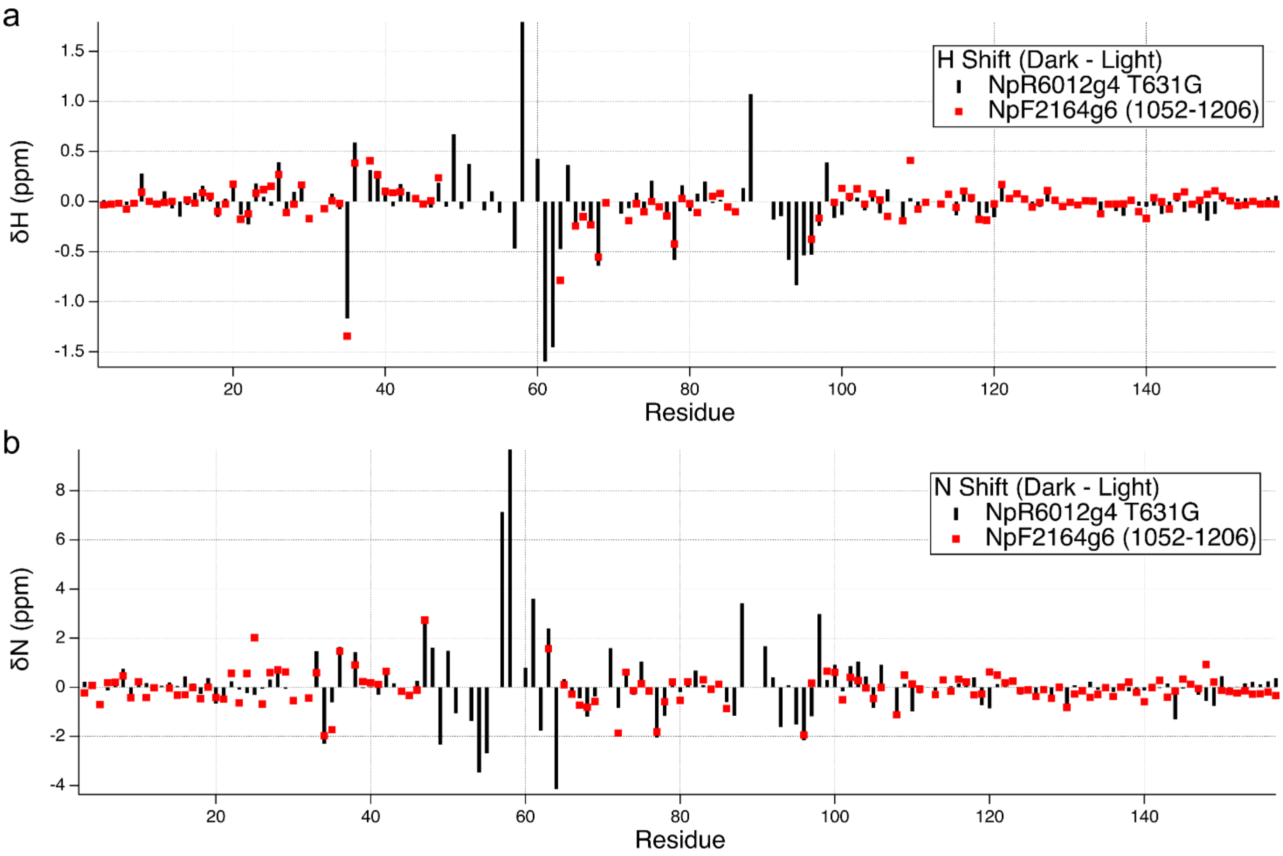


**Figure S11.** Change in chemical shift upon photoswitching. Amide hydrogen (a) and amide nitrogen (b) chemical shift changes are shown for NpR6012g4 T631G and NpF2164g6 (1052-1206). The T631G mutant of NpR6012g4 was assigned by transfer of assignments from NpR6012g4 as described previously (6). The proteins are highly homologous (Fig. S9) though they differ markedly in half-life (6).

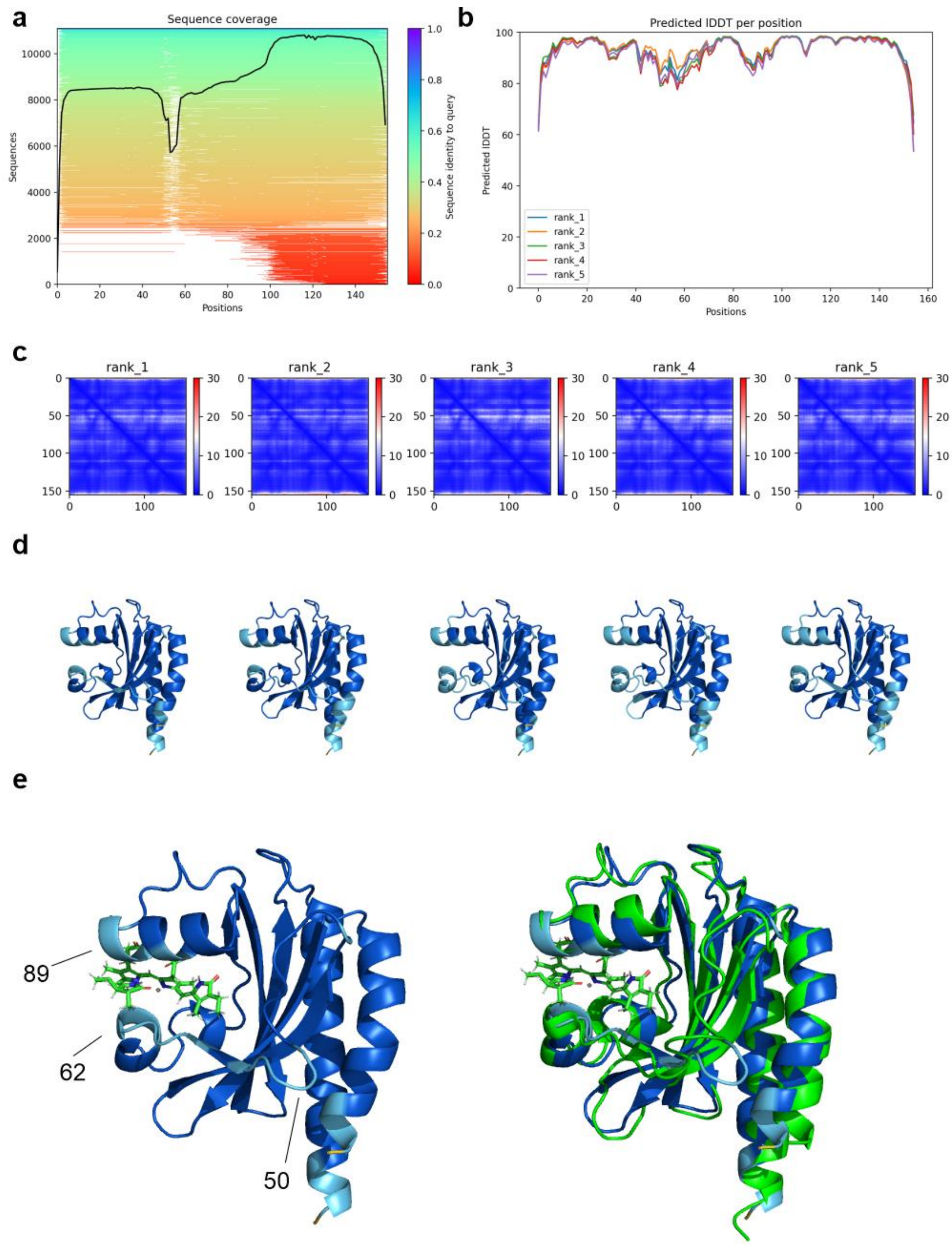


**Figure S12.** A model of the dark state of NpF2164g6 created using ColabFold with default parameters.[1] Predicted lDDT scores are above 90 for most of the sequence except for the extended loop (residues 50-62; plDDT ~80-90), the chromophore attachment site (C89), and the extreme N- and C-termini. The top-ranked model coloured by pLDDT with the PCB chromophore added and residue numbers indicated at sites of lower pLDDT. (right) An overlay of the top-ranked model, with the experimentally determined structure of NpR6012g4 (green).

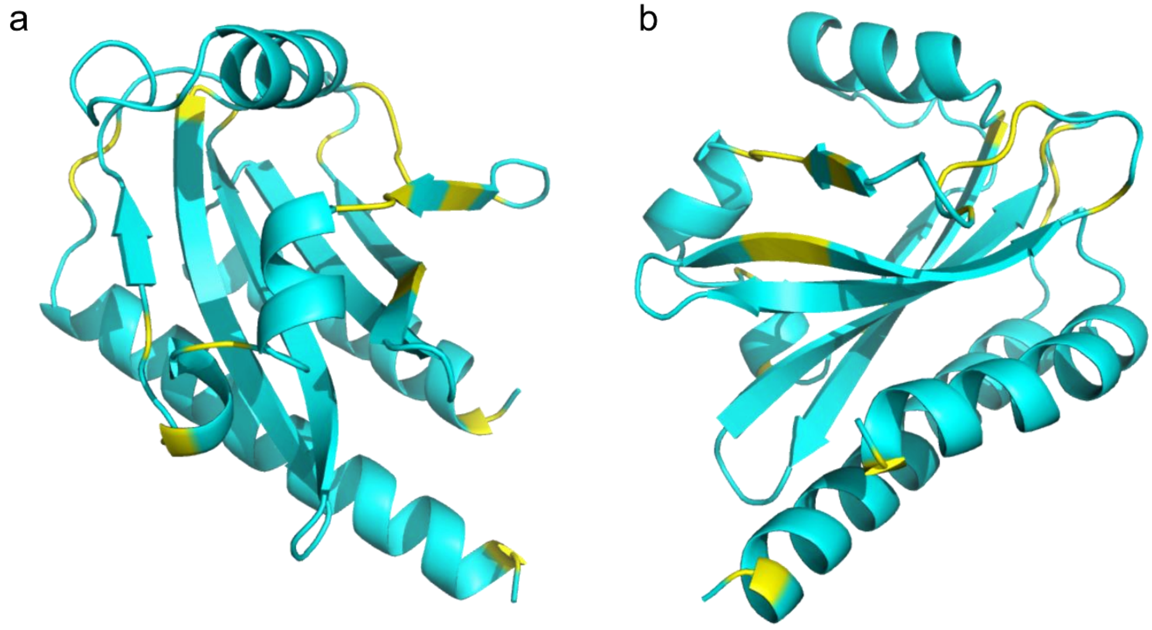

**Figure S13.** Comparison of AlphaFold[2] secondary structure and TALOSn[3] predicted secondary structure from two angles (a) and (b). Cyan residues agree in secondary structure category from TALOSn and AlphaFold (as determined by the secstruct function of CPPTRAJ[4], while yellow residues are those for which TALOSn and AlphaFold disagree.

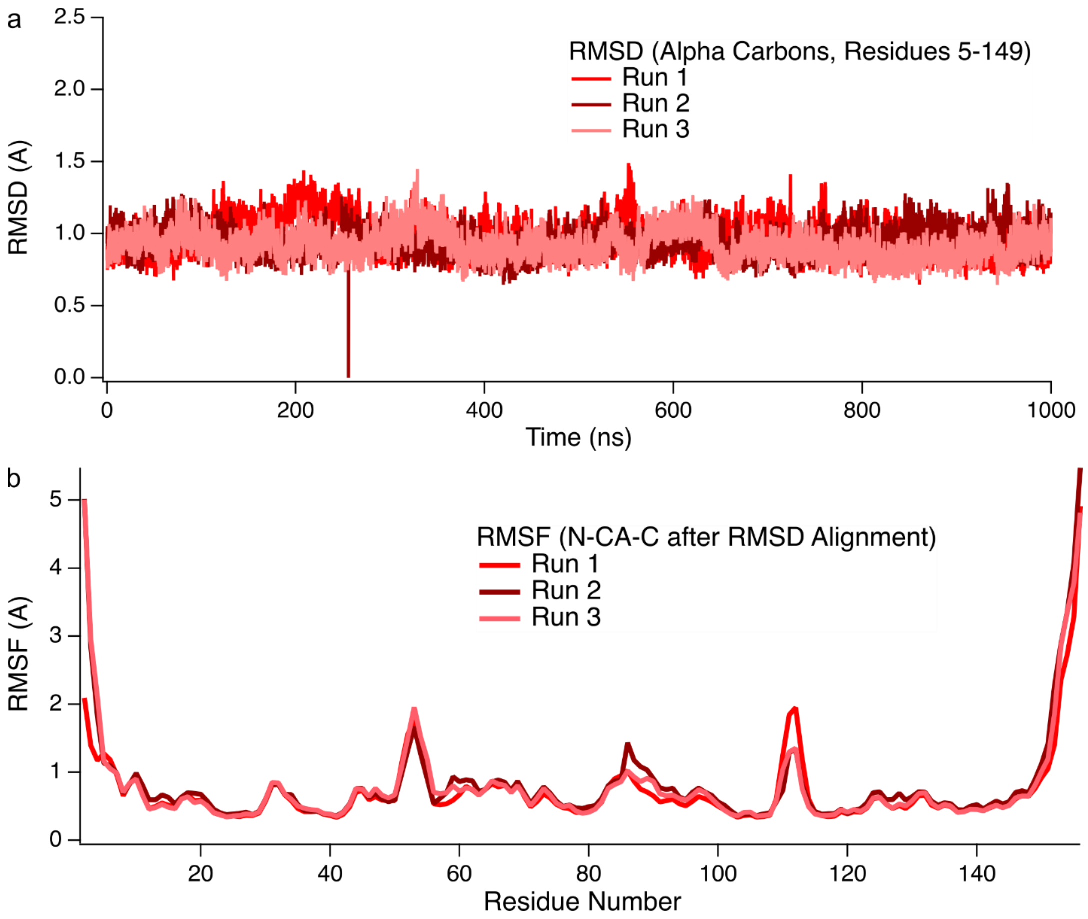


**Figure S14.** (a) The calculated α-carbon RMSD was stable at ~1Å over the course of these trajectories confirming the model represented a stable structure.[5] (b) Root mean squared fluctuations (RMSF) indicate flexibility at the termini and some of the loops in the protein, but overall stability.

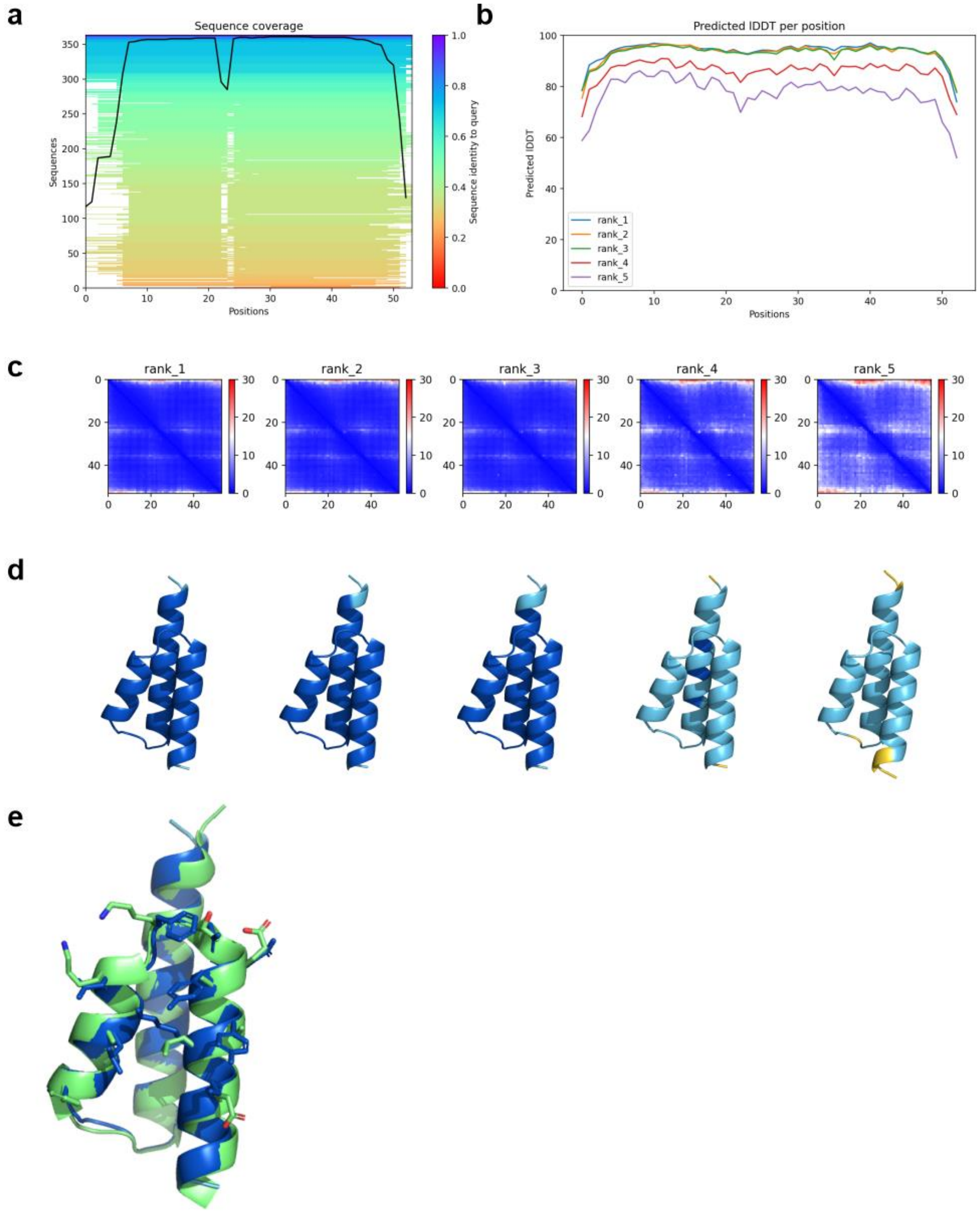


**Figure S15.** We used ColabFold[1] to generate a structure of BNp-Red-1.0 for input to the program. This structure was essentially the same as the wild-type GA domain (PDB code:1TF0, green) as expected. (a) Multiple sequence alignment (MSA) depth across the BNp-Red-1.0 sequence. (b) Predicted lDDT versus position for the 5 models generated. (c) Positional alignment error (PAE) presented as a residue-versus-residue matrix for the 5 models. (d) The 5 models coloured by pLDDT. (e) Overlay of the BNp-Red-1.0 top-ranked model (blue) and the wild-type GA domain (1TF0, green).

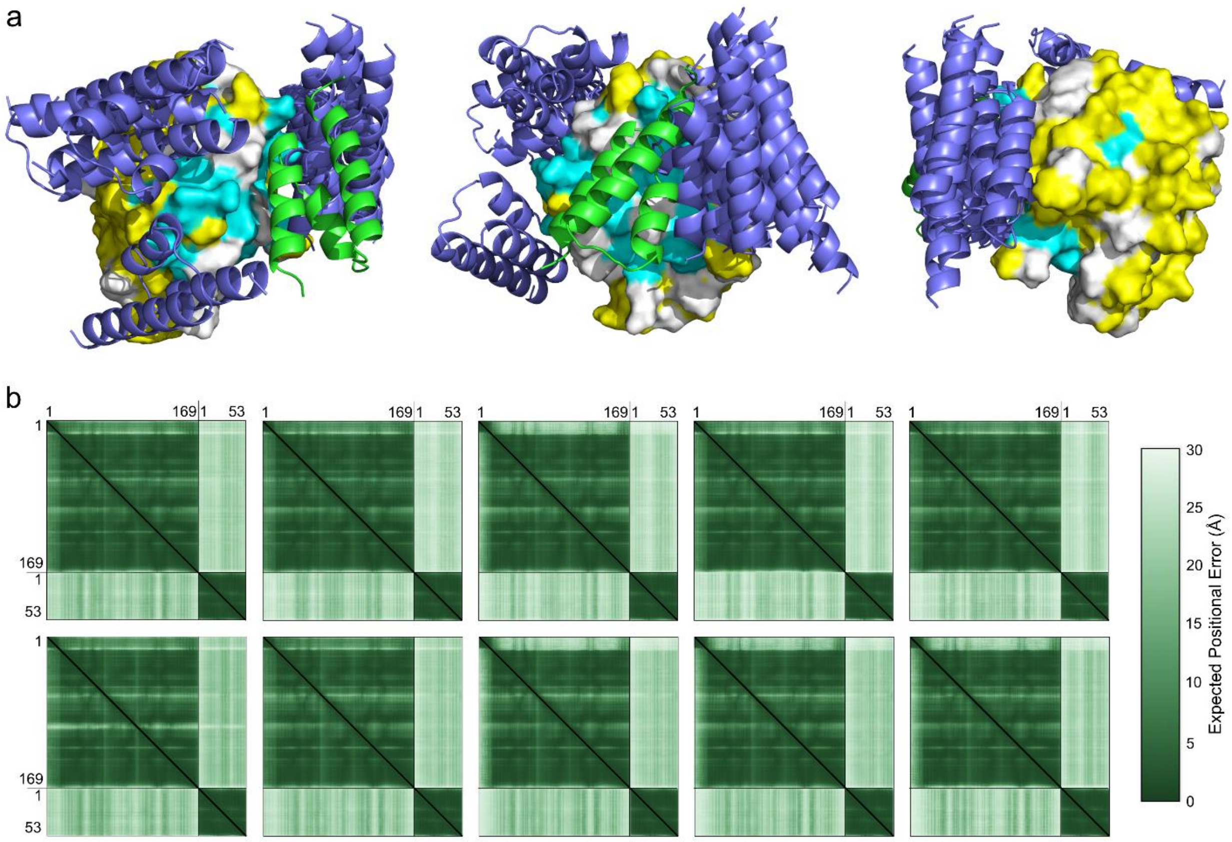


**Figure S16.** AlphaFold3 predictions for the complex of NpF2164g6 with the binders. AlphaFold3[2] was used with the sequences of NpF2164g6 1038-1206 and BNp-Red-1.2 (the tightest binder).

(a) The structures of all five models from two runs were aligned by NpF2164g6, then overlaid with the model of NpF2164g6 1054-1206 (shown as a surface) and BNp-Red-1.0 (shown as a green cartoon) generated by our combined NMR and MD study. All BNp-Red-1.2 models are depicted as purple cartoons, and the full-length NpF2164g6 is not shown for visual clarity. On the surface model of truncated NpF2164g6, residues that interact with BNp-Red-1.0 (based on NMR data) are coloured cyan and those unaffected by BNp-Red-1.0 addition are coloured yellow. Residues for which there is no data are coloured white. Three views are shown to better illustrate the structures.

(b) Residue-versus-residue predicted aligned error (PAE), is shown for the five replicates (models 0 to 4 shown from left to right) and two runs (run 1 in the top row, run 2 in the bottom row). The position and orientation of BNp-Red-1.2 as predicted by AlphaFold3 is highly variable, and not in good agreement with the NMR data. This is clear both from the variability in BNp-Red-1.2 position and orientation in (a) and the high PAE between the NpF2164g6 and BNp-Red-1.2 residues in (b). This emphasizes the difficulty AlphaFold3 has in predicting structural complexes that have no natural homologs.

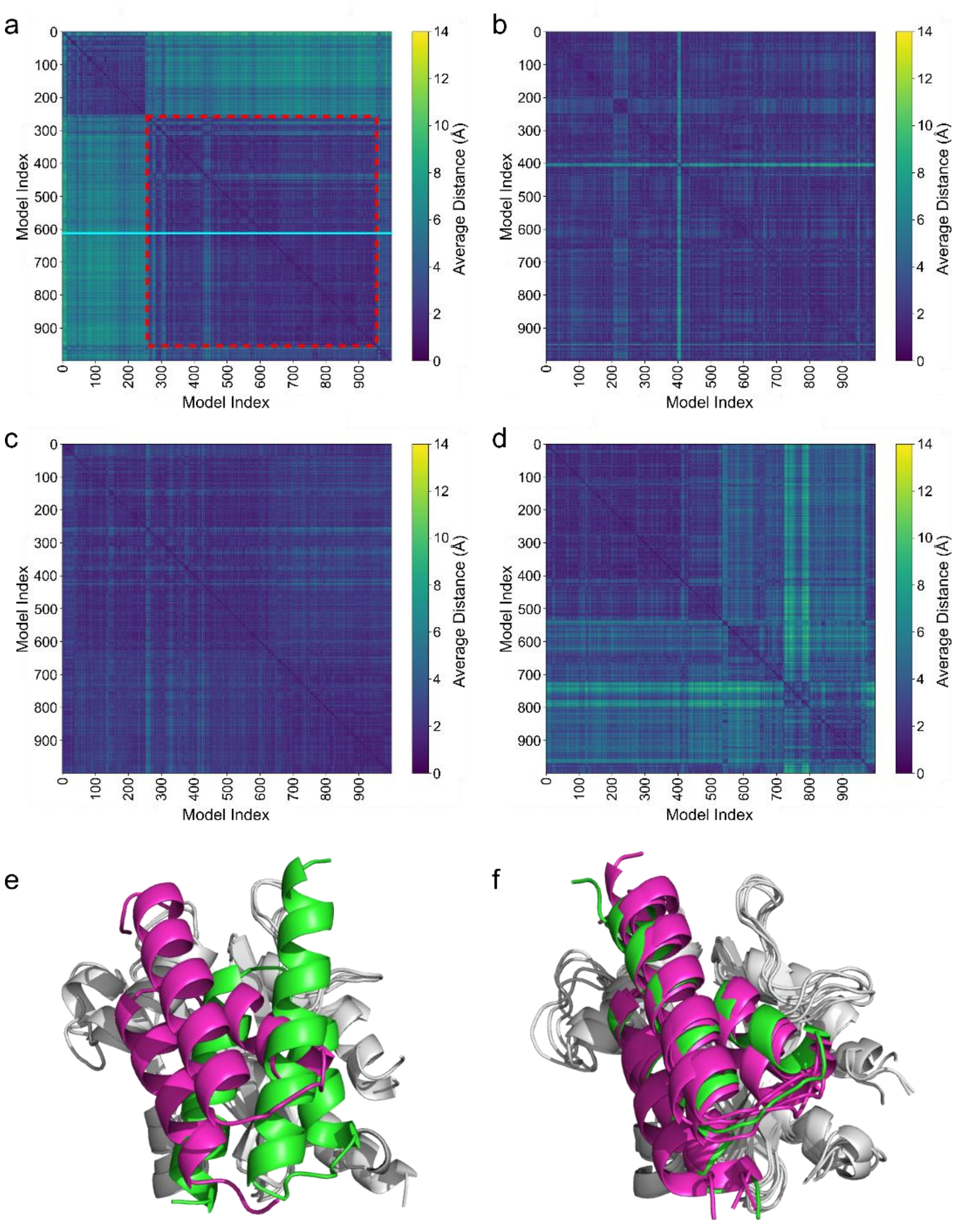


**Figure S17**. Assessment of the stability of the top-ranked structure of HADDOCK2.4 cluster 1, which was the largest in terms of number of structures (106 of 200 structures) and the second-top-ranked by the HADDOCK energy function (see Table S1 for complete statistics) (a) The MD simulation frame-versus-frame displacement metric described in the main text (see Methods) for this structure. A stable region was identified (red dashed box) from 250 ns to 950 ns, and a representative frame was determined by summation of the similarity metric across the 250 ns – 950 ns window. This frame, from the 611 ns time point, is indicated by a horizontal cyan line. (b)-(d) MD simulation frame-versus-frame displacement metric for replicates 1 to 3 that used frame 611 from (a) as a starting point. While the simulations of cluster 1 frame 611 are not entirely homogeneous, no large-scale persistent change in binder orientation is present in these trajectories. (e) The initial (green) and final (magenta) orientations of BNp-Red-1.0 from the simulation in (a) NpF2164g6 is shown in white. (f) The initial (green) and final (magenta) orientations of BNp-Red-1.0 in the three simulations of HADDOCK cluster 1 frame 611 shown in (b,c,d).

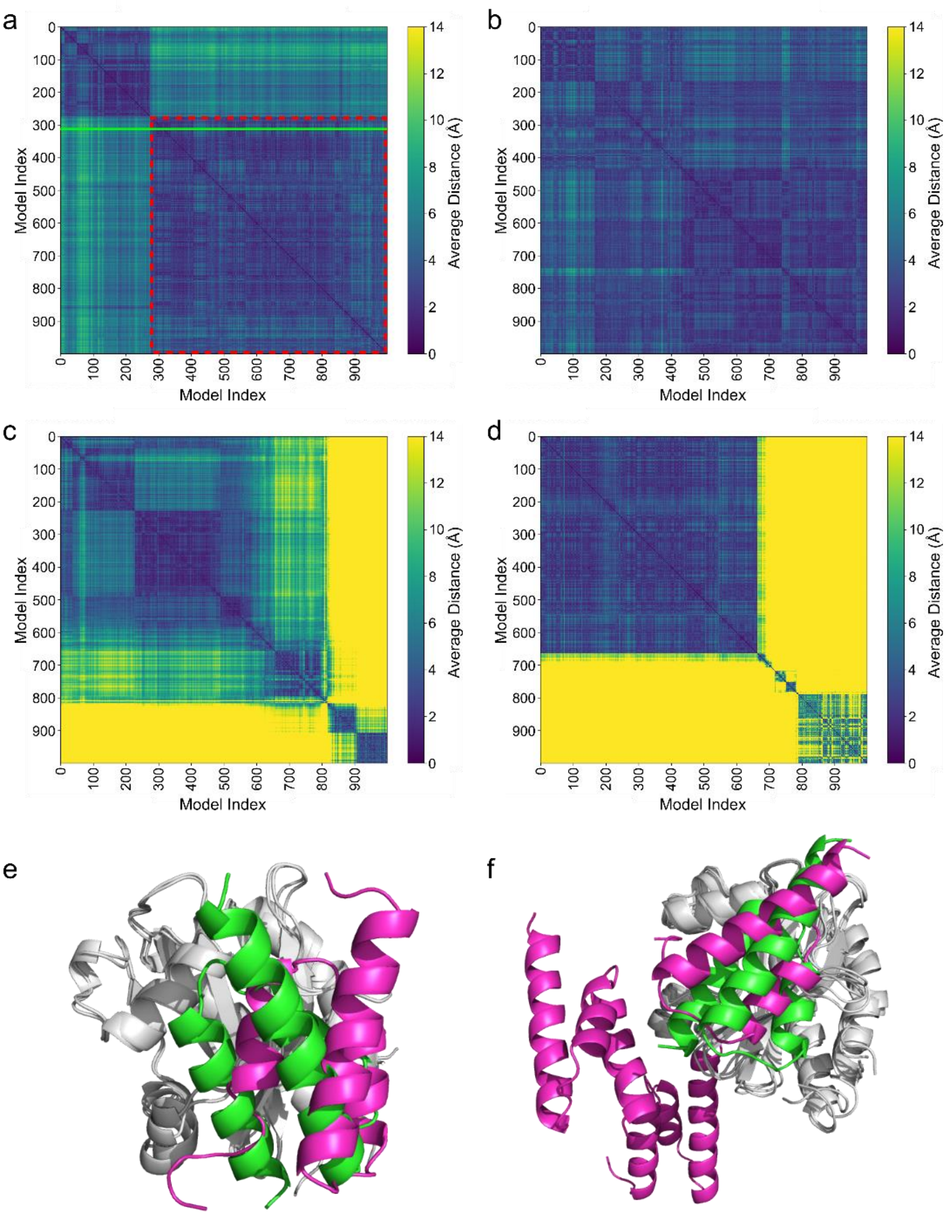


**Figure S18**. Assessment of the stability of the top-ranked structure of HADDOCK2.4 cluster 2, which was the second largest in terms of number of structures (54 of 200 structures) and the top-ranked by the HADDOCK energy function (see Table S1 for complete statistics) (a) The MD simulation frame-versus-frame displacement metric described for this structure. A relatively stable region was identified (red dashed box) from 280 ns to 1000 ns, and a representative frame was determined by summation of the similarity metric across the 280 ns – 100 ns window. The frame, from the 310 ns time point, is indicated by a horizontal green line. (b)-(d) MD simulation frame-versus-frame displacement metric for replicates 1 to 3 that used frame 310 from (a) as a starting point. (e) The initial (green) and final (magenta) orientations of BNp-Red-1.0 from the simulation in (a). NpF2164g6 is shown in white. (f) The initial (green) and final (magenta) orientations of BNp-Red-1.0 in the three simulations of HADDOCK cluster 2 frame 310 shown in (b,c,d). The complex falls apart in 2 of the 3 replicates, and in the third still undergoes a significant degree of motion. This indicates that cluster 2 is a worse candidate for the structural complex than the cluster 1.

| **Metric** | **Cluster 1** | **Cluster 2** |
|---|---|---|
| HADDOCK score (arbitrary units) | -84.9 ± 2.7 | -93.0 ± 4.4 |
| Cluster size (out of 200) | 106 | 54 |
| RMSD from the overall lowest-energy structure (Å) | 4.1 ± 0.3 | 0.6 ± 0.4 |
| Van der Waals energy (kcal/mol) | -22.1 ± 12.2 | -36.0 ± 4.5 |
| Electrostatic energy (kcal/mol) | -387.9 ± 79.2 | -374.6 ± 47.0 |
| Desolvation energy (kcal/mol) | -4.8 ± 4.9 | 2.3 ± 3.5 |
| Restraints violation energy (kcal/mol) | 196.1 ± 63.2 | 155.5 ± 43.3 |
| Buried surface area ($Å^2$) | 1260.0 ± 96.6 | 1428.2 ± 65.1 |
| Z-score | -0.7 | -1.4 |

**Table S1.** Metrics for the top two clusters of the BNp-Red-1.0/NpF2164g6 complex produced by HADDOCK2.4. The clusters are collections of docking orientations with similar contacts and inter-model RMSD, and the metrics are calculated from the 4 best-ranked structures for each cluster.

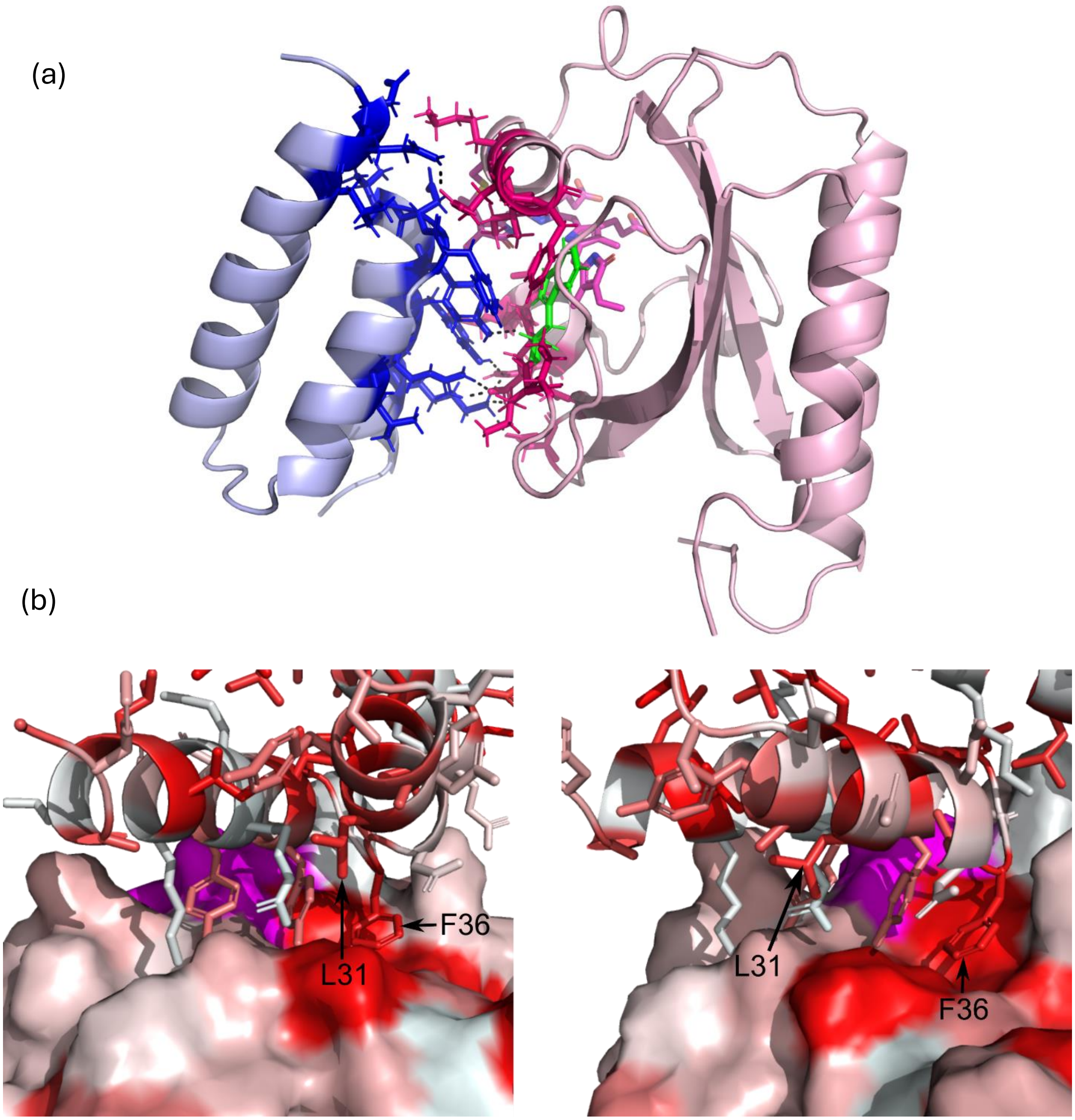


**Figure S19.** (a) The Prodigy webserver,[6] which calculates the number of interfacial contacts as well as the properties of the non-interacting surfaces,[7] identifies 30 distinct residue-residue contacts and predicts a binding affinity of -7.7 kcal.mol$^{-1}$ and a dissociation constant at 25 ºC of 2.6 μM. (b) The cluster 1 time point 611 ns snapshot, with NpF2164g6 shown as a surface and BNp-Red-1.0 shown as a cartoon with sidechains as sticks. Residues as coloured by hydrophobicity, with red being more hydrophobic and whiter being more hydrophilic. The chromophore is coloured magenta. The Phe36 residue of BNp-Red-1.0 is situated in a highly hydrophobic cleft near the chromophore. This position can mutate, but only to other hydrophobic residues (see Figure S2). Residue Leu31 of BNp-Red-1.0 is also near this pocket. In binder BNp-Red-1.2 this residue is mutated to tryptophan. This may result in additional hydrophobic interactions and explain the greater affinity of BNp-Red-1.2 compared to BNp-Red-1.0.

**Supplementary Table 2.** Plasmids designed and used in this study

| Plasmid name | Plasmid vector | Expression in | Promoter | Features | Used in | Reference |
|---|---|---|---|---|---|---|
| P1 | pBAD/HisB | Bacteria | araBAD | Avi-TEV-NpF2164g6-His6 | Figure 1c-d | This work |
| P2 | pBAD/HisB | Bacteria | araBAD | Avi-NpF2164g6-His6 | Figure 1e, S2 | This work |
| P3 | pPL | Bacteria | lac/ara-1 | PCB biosynthetic plasmid | - | Gambetta, G. *et al.* 2001 [8] |
| P4 | pMH1105 | Bacteria | T7 | NpF2164g6-His6 | Figure S1, S3-4, S8 | This work |
| P5 | pMH1105 | Bacteria | T7 | NpF2164g6 (truncated)-His6 | Figure 3, 4e, S8-11 | This work |
| P6 | p8 | Bacteria | TAC | stII (secretion signal)-FLAG-GA library-truncated P8 | - | Reis, J. *et al.* 2018 [9] |
| P7 | p3 | Bacteria | TAC | stII (secretion signal)-FLAG-GA library-DKTHTCGRP (dimerization sequence)-Cterm. P3 | Figure S2 | Reis, J. *et al.* 2018 [9] |
| P8 | pET24b | Bacteria | T7 | BNp-Red-1.0-His6 | Figure 1b, 1d, 1f, 3, 4e, S3-4, S9 | This work |
| P9 | pET24b | Bacteria | T7 | BNp-Red-1.1-His6 | Figure 1b | This work |
| P10 | pET24b | Bacteria | T7 | BNp-Red-1.2-His6 | Figure 1b, 1f, S3-4 | This work |
| P11 | pET24b | Bacteria | T7 | FLAG-BNp-Red-1.1-His6 | Figure 1e | This work |
| P12 | pET24b | Bacteria | T7 | FLAG-BNp-Red-1.2-His6 | Figure 1e | This work |
| P13 | pET24b | Bacteria | T7 | mCherry-BNp-Red-1.2-His6 | Figure 2b | This work |
| P14 | pET24b | Bacteria | T7 | FLAG-BNp-Red-1.2-mCherry-His6 | Figure S5 | This work |
| P15 | SB100X | Mammalian | EF1a ::RPBSA | ITR-$P_{EF1a}$-NpF2164g6-FUS-VP16-NLS-IRES-E-BNp-Red-1.2-NLS-pA ::$P_{RPBSA}$-EGFP-P2A-Puro-pA-ITR | Figure 2d | This work |
| P16 | SB100X | Mammalian | CMVmin ::RPBSA | ITR-etr8-$P_{CMVmin}$-SEAP-pA::$P_{RPBSA}$-HygR-pA-ITR | Figure 2d | Beyer, H. *et al.* 2024 [10] |

**Supplementary Note.** DNA sequences of plasmids designed and used in this study.

P1 Avi-TEV-NpF2164g6-His6

ATGGGTCTGAACGACATCTTCGAGGCTCAGAAAATCGAATGGCACGAAGAGGATCTGTACTTT
CAGAGCGCGGCCGAACGAGCATTAACACGGCTGTCAGAAAAAATTCGCCAAGTTCAAGAGC
TAGATACAATTTTCCGCAATGCCCTACCAGAATTGCGATCGCATTTGGAATGCGATCGCCTAGC
CGTATATCGCTTCAATCCAGATTGGGGTGGGGAGTTTATTGCCGAATCCGTAAGTCGTGAATGG
GTGGCTTTGGTTGGTCCCGAGATCAGGACTATATGGGAAGACGAACACCTACAACAAACCCA
AGGTGGACGCTATCGCAACAACGAGACGTTTGTGATTAATGACGTTTACACTGCTGGCCATGC
CCAGTGCCACTTGAAAATTCTCGAGCAATTTCAGATTAGAGCCTATATCATCACCCCGATTATT
GCCGGAAACAAGCTCTGGGGTTTGTTAGGAGCATATCAGAATAGCGGGTCGCGGCAATGGCA
AGAGAATGAAGTCAACTTGGTAGCGAAAATCGGCACGCAGTTTGGGGTTGCCGTCCAACAAT
CGCAATACTTACAACAAGGGAGCTCTGGACTCGAGCACCACCACCACCACCACTGA

P2 Avi-NpF2164g6-His6

ATGGGTCTGAACGACATCTTCGAGGCTCAGAAAATCGAATGGCACGAAGGGTCAAGCGGAG
AACGAGCATTAACACGGCTGTCAGAAAAAATTCGCCAAGTTCAAGAGCTAGATACAATTTTCC
GCAATGCCCTACCAGAATTGCGATCGCATTTGGAATGCGATCGCCTAGCCGTATATCGCTTCAA
TCCAGATTGGGGTGGGGAGTTTATTGCCGAATCCGTAAGTCGTGAATGGGTGGCTTTGGTTGG
TCCCGAGATCAGGACTATATGGGAAGACGAACACCTACAACAAACCCAAGGTGGACGCTATC
GCAACAACGAGACGTTTGTGATTAATGACGTTTACACTGCTGGCCATGCCCAGTGCCACTTGA
AAATTCTCGAGCAATTTCAGATTAGAGCCTATATCATCACCCCGATTATTGCCGGAAACAAGCT
CTGGGGTTTGTTAGGAGCATATCAGAATAGCGGGTCGCGGCAATGGCAAGAGAATGAAGTCA
ACTTGGTAGCGAAAATCGGCACGCAGTTTGGGGTTGCCGTCCAACAATCGCAATACTTACAAC
AAGGGAGCTCTGGACTCGAGCACCACCACCACCACCACTGA

P4 NpF2164g6-His6

ATGGAACGAGCATTAACACGGCTGTCAGAAAAAATTCGCCAAGTTCAAGAGCTAGATACAAT
TTTCCGCAATGCCCTACCAGAATTGCGATCGCATTTGGAATGCGATCGCCTAGCCGTATATCGC
TTCAATCCAGATTGGGGTGGGGAGTTTATTGCCGAATCCGTAAGTCGTGAATGGGTGGCTTTG
GTTGGTCCCGAGATCAGGACTATATGGGAAGACGAACACCTACAACAAACCCAAGGTGGACG
CTATCGCAACAACGAGACGTTTGTGATTAATGACGTTTACACTGCTGGCCATGCCCAGTGCCA
CTTGAAAATTCTCGAGCAATTTCAGATTAGAGCCTATATCATCACCCCGATTATTGCCGGAAAC
AAGCTCTGGGGTTTGTTAGGAGCATATCAGAATAGCGGGTCGCGGCAATGGCAAGAGAATGA
AGTCAACTTGGTAGCGAAAATCGGCACGCAGTTTGGGGTTGCCGTCCAACAATCGCAATACTT
ACAACAAGGGAGCTCTGGACATCATCACCATCACCATTAA

P5 NpF2164g6 (truncated)-His6

ATGCAAGAGCTAGATACAATTTTCCGCAATGCCCTACCAGAATTGCGATCGCATTTGGAATGCG
ATCGCCTAGCCGTATATCGCTTCAATCCAGATTGGGGTGGGGAGTTTATTGCCGAATCCGTAAG
TCGTGAATGGGTGGCTTTGGTTGGTCCCGAGATCAGGACTATATGGGAAGACGAACACCTACA
ACAAACCCAAGGTGGACGCTATCGCAACAACGAGACGTTTGTGATTAATGACGTTTACACTGC
TGGCCATGCCCAGTGCCACTTGAAAATTCTCGAGCAATTTCAGATTAGAGCCTATATCATCACC
CCGATTATTGCCGGAAACAAGCTCTGGGGTTTGTTAGGAGCATATCAGAATAGCGGGTCGCGG

CAATGGCAAGAGAATGAAGTCAACTTGGTAGCGAAAATCGGCACGCAGTTTGGGGTTGCCGT
CCAACAATCGCAATACTTACAACAAGGGAGCTCTGGACATCATCACCATCACCATTAA

P8 BNp-Red-1.0-His6

ATGAAACTGGCCACGATTGACCAGTGGCTGCTGAAAAACGCGAAAGAAGATGCTATTGCAGA
ACTGAAAAAGGCTGGTATCACCTCTGACGCGTATTTCAACTTGATCAATGATGCGTTTGATGTG
GATTATGTTAACTATCGGAAGAACAAGATCCTGAAAGCTCACGCCGGGAGCTCTGGACTCGA
GCACCACCACCACCACCACTGA

P9 BNp-Red-1.1-His6

ATGAAACTGGCCACGATTGACCAGTGGCTGCTGAAAAACGCGAAAGAAGATGCTATTGCAGA
ACTGAAAAAGGCTGGTATCACCGCTGACGCGTATTTCAACTTGATCAATGATGCGTGGGATGT
GGACTATGTTAACTATCGTAAGAACAAGATCCTGAAAGCTCACGCCGGGAGCTCTGGACTCG
AGCACCACCACCACCACCACTGA

P10 BNp-Red-1.2-His6

ATGAAACTGGCCACGATTGACCAGTGGCTGCTGAAAAACGCGAAAGAAGATGCTATTGCAGA
ACTGAAAAAGGCTGGTATCACCGCTGACGCGTATTTCAACTGGATCAATAATGCGTGGGATGT
GGATTATGTTAACTATCGGAAGAACAAGATCCTGAAAGCTCACGCCGGGAGCTCTGGACTCG
AGCACCACCACCACCACCACTGA

P11 FLAG-BNp-Red-1.1-His6

ATGAAACTGGCCGATTATAAAGATGATGATGATAAAGGCGGTTCAACGATTGACCAGTGGCTG
CTGAAAAACGCGAAAGAAGATGCTATTGCAGAACTGAAAAAGGCTGGTATCACCGCTGACGC
GTATTTCAACTTGATCAATGATGCGTGGGATGTGGACTATGTTAACTATCGTAAGAACAAGATC
CTGAAAGCTCACGCCGGGAGCTCTGGACTCGAGCACCACCACCACCACCACTGA

P12 FLAG-BNp-Red-1.2-His6

ATGAAACTGGCCGATTATAAAGATGATGATGATAAAGGCGGTTCAACGATTGACCAGTGGCTG
CTGAAAAACGCGAAAGAAGATGCTATTGCAGAACTGAAAAAGGCTGGTATCACCGCTGACGC
GTATTTCAACTGGATCAATAATGCGTGGGATGTGGATTATGTTAACTATCGGAAGAACAAGATC
CTGAAAGCTCACGCCGGGAGCTCTGGACTCGAGCACCACCACCACCACCACTGA

P13 mCherry-BNp-Red-1.2-His6

ATGGTGAGCAAGGGCGAGGAGGATAACATGGCCATCATCAAGGAGTTCATGCGCTTCAAGGT
GCACATGGAGGGCTCCGTGAACGGCCACGAGTTCGAGATCGAGGGCGAGGGCGAGGGCCGC
CCCTACGAGGGCACCCAGACCGCCAAGCTGAAGGTGACCAAGGGTGGCCCCCTGCCCTTCGC
CTGGGACATCCTGTCCCCTCAGTTCATGTACGGCTCCAAGGCCTACGTGAAGCACCCCGCCGA
CATCCCCGACTACTTGAAGCTGTCCTTCCCCGAGGGCTTCAAGTGGGAGCGCGTGATGAACTT
CGAGGACGGCGGCGTGGTGACCGTGACCCAGGACTCCTCCCTGCAGGACGGCGAGTTCATCT
ACAAGGTGAAGCTGCGCGGCACCAACTTCCCCTCCGACGGCCCTGTAATGCAGAAGAAGACT
ATGGGCTGGGAGGCCTCCTCCGAGCGGATGTACCCCGAGGACGGCGCCCTGAAGGGCGAGAT
CAAGCAGAGGCTGAAGCTGAAGGACGGCGGCCACTACGACGCTGAGGTCAAGACCACCTAC
AAGGCCAAGAAGCCCGTGCAGCTGCCCGGCGCCTACAACGTCAACATCAAGTTGGACATCAC

CTCCCACAACGAGGACTACACCATCGTGGAACAGTACGAACGCGCCGAGGGCCGCCACTCCA
CCGGCGGCATGGACGAGCTGTACAAGGGCGGATCCACGATTGACCAGTGGCTGCTGAAAAAC
GCGAAAGAAGATGCTATTGCAGAACTGAAAAAGGCTGGTATCACCGCTGACGCGTATTTCAA
CTGGATCAATAATGCGTGGGATGTGGATTATGTTAACTATCGGAAGAACAAGATCCTGAAAGC
TCACGCCGGGAGCTCTGGAAAGCTTCACCACCACCACCACCACTGA

P14 FLAG-BNp-Red-1.2-mCherry-His6

ATGAAACTGGCCGATTATAAAGATGATGATGATAAAGGCGGTTCAACGATTGACCAGTGGCTG
CTGAAAAACGCGAAAGAAGATGCTATTGCAGAACTGAAAAAGGCTGGTATCACCGCTGACGC
GTATTTCAACTGGATCAATAATGCGTGGGATGTGGATTATGTTAACTATCGGAAGAACAAGATC
CTGAAAGCTCACGCCGGGAGCTCTGGAGTGAGCAAGGGCGAGGAGGATAACATGGCCATCAT
CAAGGAGTTCATGCGCTTCAAGGTGCACATGGAGGGCTCCGTGAACGGCCACGAGTTCGAGA
TCGAGGGCGAGGGCGAGGGCCGCCCCTACGAGGGCACCCAGACCGCCAAGCTGAAGGTGAC
CAAGGGTGGCCCCCTGCCCTTCGCCTGGGACATCCTGTCCCCTCAGTTCATGTACGGCTCCAA
GGCCTACGTGAAGCACCCCGCCGACATCCCCGACTACTTGAAGCTGTCCTTCCCCGAGGGCTT
CAAGTGGGAGCGCGTGATGAACTTCGAGGACGGCGGCGTGGTGACCGTGACCCAGGACTCC
TCCCTGCAGGACGGCGAGTTCATCTACAAGGTGAAGCTGCGCGGCACCAACTTCCCCTCCGA
CGGCCCTGTAATGCAGAAGAAGACTATGGGCTGGGAGGCCTCCTCCGAGCGGATGTACCCCG
AGGACGGCGCCCTGAAGGGCGAGATCAAGCAGAGGCTGAAGCTGAAGGACGGCGGCCACTA
CGACGCTGAGGTCAAGACCACCTACAAGGCCAAGAAGCCCGTGCAGCTGCCCGGCGCCTAC
AACGTCAACATCAAGTTGGACATCACCTCCCACAACGAGGACTACACCATCGTGGAACAGTA
CGAACGCGCCGAGGGCCGCCACTCCACCGGCGGCATGGACGAGCTGTACAAGCTCGAGCAC
CACCACCACCACCACTGA

P15-1 NpF2164g6-FUS-VP16-NLS-IRES-E-BNp-Red-1.2-NLS

ATGGAACGAGCATTAACACGGCTGTCAGAAAAAATTCGCCAAGTTCAAGAGCTAGATACAAT
TTTCCGCAATGCCCTACCAGAATTGCGATCGCATTTGGAATGCGATCGCCTAGCCGTATATCGC
TTCAATCCAGATTGGGGTGGGGAGTTTATTGCCGAATCCGTAAGTCGTGAATGGGTGGCTTTG
GTTGGTCCCGAGATCAGGACTATATGGGAAGACGAACACCTACAACAAACCCAAGGTGGACG
CTATCGCAACAACGAGACGTTTGTGATTAATGACGTTTACACTGCTGGCCATGCCCAGTGCCA
CTTGAAAATTCTCGAGCAATTTCAGATTAGAGCCTATATCATCACCCCGATTATTGCCGGAAAC
AAGCTCTGGGGTTTGTTAGGAGCATATCAGAATAGCGGGTCGCGGCAATGGCAAGAGAATGA
AGTCAACTTGGTAGCGAAAATCGGCACGCAGTTTGGGGTTGCCGTCCAACAATCGCAATACTT
ACAACAAGAATTCATGGCCTCAAACGATTATACCCAACAAGCAACCCAAAGCTATGGGGCCTA
CCCCACCCAGCCCGGGCAGGGCTATTCCCAGCAGAGCAGTCAGCCCTACGGACAGCAGAGTT
ACAGTGGTTATAGCCAGTCCACGGACACTTCAGGCTATGGCCAGAGCAGCTATTCTTCTTATG
GCCAGAGCCAGAACACAGGCTATGGAACTCAGTCAACTCCCCAGGGATATGGCTCGACTGGC
GGCTATGGCAGTAGCCAGAGCTCCCAATCGTCTTACGGGCAGCAGTCCTCCTACCCTGGCTAT
GGCCAGCAGCCAGCTCCCAGCAGCACCTCGGGAAGTTACGGTAGCAGTTCTCAGAGCAGCA
GCTATGGGCAGCCCCAGAGTGGGAGCTACAGCCAACAGCCTAGCTATGGTGGACAGCAGCAA
TCTTACGGTCAACAACAGAGCTATAATCCCCCTCAGGGCTATGGACAGCAGAACCAGTACAAC
AGCAGCAGTGGTGGTGGAGGTGGAGGTGGAGGTGGAGGTAACTATGGCCAAGATCAATCCTC
CATGAGTAGTGGTGGTGGCAGTGGTGGCGGTTATGGCAATCAAGACCAGAGTGGTGGAGGTG
GCAGCGGTGGCTATGGACAGCAGGACCGTGGATCCGCGTACAGCCGCGCGCGTACGAAAAAC

AATTACGGGTCTACCATCGAGGGCCTGCTCGATCTCCCGGACGACGACGCCCCCGAAGAGGC
GGGGCTGGCGGCTCCGCGCCTGTCCTTTCTCCCCGCGGGACACACGCGCAGACTGTCGACGG
CCCCCCCGACCGATGTCAGCCTGGGGGACGAGCTCCACTTAGACGGCGAGGACGTGGCGATG
GCGCATGCCGACGCGCTAGACGATTTCGATCTGGACATGTTGGGGGACGGGGATTCCCCGGGT
CCGGGATTTACCCCCCACGACTCCGCCCCCTACGGCGCTCTGGATATGGCCGACTTCGAGTTT
GAGCAGATGTTTACCGATGCCCTTGGAATTGACGAGTACGGTGGGCCCAAGAAAAAGCGGAA
GGTGTGATCTAGAGTCGACCTGCAGCCCAAGCTTAAAACAGCTCTGGGGTTGTACCCACCCC
AGAGGCCCACGTGGCGGCTAGTACTCCGGTATTGCGGTACCCTTGTACGCCTGTTTTATACTCC
CTTCCCGTAACTTAGACGCACAAAACCAAGTTCAATAGAAGGGGGTACAAACCAGTACCACC
ACGAACAAGCACTTCTGTTTCCCCGGTGATGTCGTATAGACTGCTTGCGTGGTTGAAAGCGAC
GGATCCGTTATCCGCTTATGTACTTCGAGAAGCCCAGTACCACCTCGGAATCTTCGATGCGTTG
CGCTCAGCACTCAACCCCAGAGTGTAGCTTAGGCTGATGAGTCTGGACATCCCTCACCGGTG
ACGGTGGTCCAGGCTGCGTTGGCGGCCTACCTATGGCTAACGCCATGGGACGCTAGTTGTGAA
CAAGGTGTGAAGAGCCTATTGAGCTACATAAGAATCCTCCGGCCCCTGAATGCGGCTAATCCC
AACCTCGGAGCAGGTGGTCACAAACCAGTGATTGGCCTGTCGTAACGCGCAAGTCCGTGGCG
GAACCGACTACTTTGGGTGTCCGTGTTTCCTTTTATTTTATTGTGGCTGCTTATGGTGACAATCA
CAGATTGTTATCATAAAGCGAATTGGATTGCGGCGATATCGCCACCATGCCCCGCCCCAAGCTC
AAGTCCGATGACGAGGTACTCGAGGCCGCCACCGTAGTGCTGAAGCGTTGCGGTCCCATAGA
GTTCACGCTCAGCGGAGTAGCAAAGGAGGTGGGGCTCTCCCGCGCAGCGTTAATCCAGCGCT
TCACCAACCGCGATACGCTGCTGGTGAGGATGATGGAGCGCGGCGTCGAGCAGGTGCGGCAT
TACCTGAATGCGATACCGATAGGCGCAGGGCCGCAAGGGCTCTGGGAATTTTTGCAGGTGCTC
GTTCGGAGCATGAACACTCGCAACGACTTCTCGGTGAACTATCTCATCTCCTGGTACGAGCTC
CAGGTGCCGGAGCTACGCACGCTTGCGATCCAGCGGAACCGCGCGGTGGTGGAGGGGATCC
GCAAGCGACTGCCCCCAGGTGCTCCTGCGGCAGCTGAGTTGCTCCTGCACTCGGTCATCGCT
GGCGCGACGATGCAGTGGGCCGTCGATCCGGATGGTGAGCTAGCTGATCATGTGCTGGCTCA
GATCGCTGCCATCCTGTGTTTAATGTTTCCCGAACACGACGATTTCCAACTCCTCCAGGCACAT
GCGTCCGCGTACAGCTTAATTGTACATAACAGTGCTGGTAGTGCTGGTAGTGCTGGTATGAAA
CTGGCCACGATTGACCAGTGGCTGCTGAAAAACGCGAAAGAAGATGCTATTGCAGAACTGAA
AAAGGCTGGTATCACCGCTGACGCGTATTTCAACTGGATCAATAATGCGTGGGATGTGGATTAT
GTTAACTATCGGAAGAACAAGATCCTGAAAGCTCACGCCCCCAAGAAAAAGCGGAAGGTGT
GA

P15-2 EGFP-P2A-Puro

ATGTTGAGCAAGGGCGAGGAGCTGTTCACCGGGGTGGTGCCCATCCTGGTCGAGCTGGACGG
CGACGTAAACGGCCACAAGTTCAGCGTGTCCGGCGAGGGCGAGGGCGATGCCACCTACGGC
AAGCTGACCCTGAAGTTCATCTGCACCACCGGCAAGCTGCCCGTGCCCTGGCCCACCCTCGT
GACCACCCTGACCTACGGCGTGCAGTGCTTCAGCCGCTACCCCGACCACATGAAGCAGCACG
ACTTCTTCAAGTCCGCCATGCCCGAAGGCTACGTCCAGGAGCGCACCATCTTCTTCAAGGACG
ACGGCAACTACAAGACCCGCGCCGAGGTGAAGTTCGAGGGCGACACCCTGGTGAACCGCAT
CGAGCTGAAGGGCATCGACTTCAAGGAGGACGGCAACATCCTGGGGCACAAGCTGGAGTAC
AACTACAACAGCCACAACGTCTATATCATGGCCGACAAGCAGAAGAACGGCATCAAGGTGAA
CTTCAAGATCCGCCACAACATCGAGGACGGCAGCGTGCAGCTCGCCGACCACTACCAGCAGA
ACACCCCCATCGGCGACGGCCCCGTGCTGCTGCCCGACAACCACTACCTGAGCACCCAGTCC
GCCCTGAGCAAAGACCCCAACGAGAAGCGCGATCACATGGTCCTGCTGGAGTTCGTGACCGC

CGCCGGGATCACTCTCGGCATGGACGAGCTGTACAAGGGCAGTGGAGCTACTAACTTCAGCC
TGCTGAAGCAGGCTGGTGACGTCGAGGAGAATCCTGGCCCCATGACCGAGTACAAGCCCACG
GTGCGCCTCGCCACCCGCGACGACGTCCCCAGGGCCGTACGCACCCTCGCCGCCGCGTTCGC
CGACTACCCCGCCACGCGCCACACCGTCGATCCGGACCGCCACATCGAGCGGGTCACCGAGC
TGCAAGAACTCTTCCTCACGCGCGTCGGGCTCGACATCGGCAAGGTGTGGGTCGCGGACGAC
GGCGCCGCGGTGGCGGTCTGGACCACGCCGGAGAGCGTCGAAGCGGGGGCGGTGTTCGCCG
AGATCGGCCCGCGCATGGCCGAGTTGAGCGGTTCCCGGCTGGCCGCGCAGCAACAGATGGAA
GGCCTCCTGGCGCCGCACCGGCCCAAGGAGCCCGCGTGGTTCCTGGCCACCGTCGGCGTCTC
GCCCGACCACCAGGGCAAGGGTCTGGGCAGCGCCGTCGTGCTCCCCGGAGTGGAGGCGGCC
GAGCGCGCCGGGGTGCCCGCCTTCCTGGAGACCTCCGCGCCCCGCAACCTCCCCTTCTACGA
GCGGCTCGGCTTCACCGTCACCGCCGACGTCGAGGTGCCCGAAGGACCGCGCACCTGGTGC
ATGACCCGCAAGCCCGGTGCCTGA

#### References


1. M. Mirdita, K. Schutze, Y. Moriwaki, L. Heo, S. Ovchinnikov and M. Steinegger, ColabFold: making protein folding accessible to all, *Nat Methods*, 2022, **19**, 679-682.
2. J. Abramson, J. Adler, J. Dunger, R. Evans, T. Green, A. Pritzel, O. Ronneberger, L. Willmore, A. J. Ballard, J. Bambrick, S. W. Bodenstein, D. A. Evans, C. C. Hung, M. O'Neill, D. Reiman, K. Tunyasuvunakool, Z. Wu, A. Zemgulyte, E. Arvaniti, C. Beattie, O. Bertolli, A. Bridgland, A. Cherepanov, M. Congreve, A. I. Cowen-Rivers, A. Cowie, M. Figurnov, F. B. Fuchs, H. Gladman, R. Jain, Y. A. Khan, C. M. R. Low, K. Perlin, A. Potapenko, P. Savy, S. Singh, A. Stecula, A. Thillaisundaram, C. Tong, S. Yakneen, E. D. Zhong, M. Zielinski, A. Zidek, V. Bapst, P. Kohli, M. Jaderberg, D. Hassabis and J. M. Jumper, Accurate structure prediction of biomolecular interactions with AlphaFold 3, *Nature*, 2024, **630**, 493-500.
3. Y. Shen and A. Bax, Protein backbone and sidechain torsion angles predicted from NMR chemical shifts using artificial neural networks, *J Biomol NMR*, 2013, **56**, 227-241.
4. D. R. Roe and T. E. Cheatham, 3rd, PTRAJ and CPPTRAJ: Software for Processing and Analysis of Molecular Dynamics Trajectory Data, *J Chem Theory Comput*, 2013, **9**, 3084-3095.
5. L. Heo and M. Feig, Experimental accuracy in protein structure refinement via molecular dynamics simulations, *Proc Natl Acad Sci U S A*, 2018, **115**, 13276-13281.
6. L. C. Xue, J. P. Rodrigues, P. L. Kastritis, A. M. Bonvin and A. Vangone, PRODIGY: a web server for predicting the binding affinity of protein-protein complexes, *Bioinformatics*, 2016, **32**, 3676-3678.
7. P. L. Kastritis, J. P. Rodrigues, G. E. Folkers, R. Boelens and A. M. Bonvin, Proteins feel more than they see: fine-tuning of binding affinity by properties of the non-interacting surface, *J Mol Biol*, 2014, **426**, 2632-2652.
8. G. A. Gambetta and J. C. Lagarias, Genetic engineering of phytochrome biosynthesis in bacteria, *Proc Natl Acad Sci U S A*, 2001, **98**, 10566-10571.
9. J. M. Reis, X. Xu, S. McDonald, R. M. Woloschuk, A. S. I. Jaikaran, F. S. Vizeacoumar, G. A. Woolley and M. Uppalapati, Discovering Selective Binders for Photoswitchable Proteins Using Phage Display, *ACS Synth Biol*, 2018, **7**, 2355-2364.
10. H. M. Beyer, S. Kumar, M. Nieke, C. M. C. Diehl, K. Tang, S. Shumka, C. S. Koh, C. Fleck, J. A. Davies, M. Khammash and M. D. Zurbriggen, Genetically-stable engineered optogenetic gene switches modulate spatial cell morphogenesis in two- and three-dimensional tissue cultures, *Nat Commun*, 2024, **15**, 10470.
